# Three-dimensional ultrastructural differences between thalamic and non-thalamic recipient layers in macaque V1

**DOI:** 10.1101/2025.08.04.668334

**Authors:** Virginia Garcia-Marin, Michael J Hawken

## Abstract

Understanding the synaptic characteristics of each cortical layer is essential for elucidating the functional architecture of each brain region. In the current study, we made a detailed quantitative comparison of the synaptic structure in the predominantly input layers of primate primary visual cortex (layer 4C) and in the predominant output layer (layer 3B) using focused ion beam scanning electron microscopy (FIB/SEM). We quantified the synaptic density in each layer, classified synaptic boutons according to their number of synapses and mitochondrial content, and quantified key morphometric parameters, including bouton volume, postsynaptic density (PSD) area and morphology, volume occupied by mitochondria, and postsynaptic targets. Our results revealed that for all the layers there is a higher proportion of single-synapse boutons without mitochondria. Multisynaptic boutons containing mitochondria (MSBm+)— which likely correspond to TC terminals —were significantly more abundant in the thalamocortical recipient layers 4Cα and 4Cβ. These MSBm+ boutons were also larger, more likely to contact dendritic spines, and contained more mitochondria than other bouton categories. In contrast, layer 3B, displayed a lower prevalence of MSBm+ boutons, these boutons were smaller than those in layer 4C and made fewer synapses. These findings highlight laminar differences in bouton architecture and support the idea that TC synapses are structurally adapted to support high synaptic efficacy. Together, our data provide a detailed quantitative framework for understanding the synaptic organization of primate V1, with implications for sensory processing and cortical circuit function.

## INTRODUCTION

In cortex, one approach to elucidate the anatomical and physiological principles that underline brain function is to ask if the structure and function in one cortical area can be extrapolated to other cortical areas or to the same brain area in a different species. The canonical cortical circuit (Douglas & Martin 2004, 2007a,b) has been foundational to this approach. Although the canonical circuit motif shares many features between cortical areas within and across species (laminar structure, cell types and connectivity motifs) there are many key features that have yet to be elucidated. Some authors, however, question the homogeneity of the cortex and suggest that at least the modules must contain significant specialization to enable that different cortical areas perform distinct tasks (Weiler et al. 2008; Oviedo et al. 2010; Hooks et al. 2011, Constantinople & Bruno 2013). While there is a canonical microcircuitry, the specific connectivity and strength of connections can vary between different cortical regions and layers as a result from relatively small quantitative changes in the elements that build the circuits: neurons, glial cells, and synapses. For this reason, it is essential to determine relative proportions or densities of these different circuit elements, to identify different morphologies and types, and find out how these elements are connected.

Central to understanding connectivity is determining the density of synapses and the types of synapses. A large analysis of the entire mouse brain – and, in twenty regions of the human brain - using two classes of post-synaptic proteins PSD95 (Postsynaptic Density 95) and SAP102 (Synapse-Associated Protein 102), revealed that each brain area exhibits a unique synaptic distribution characterized by distinct puncta density, intensity, and size (Zhu et al. 2018; Curran et al. 2021). This data supports the idea that each region of the brain has a characteristic ‘synaptome signature’ that would reflect their functional diversity. This detailed analysis of the cortex signature at the level of neurons and synapses has become central in endeavors to build realistic computational models that mimic the physiological functioning of circuits (review in Vanni et al. 2020).

Structural data about cortical circuits can be obtained at different scales: microscopic level, for the neuronal quantification and nanoscopic scale for the synaptic analysis (Sporns et al. 2005; DeFelipe, 2010; Dorkenwald et al. 2024; Schlegel et al. 2024). Recent advances involving serial electron microscopy (EM) and subsequent imaging methods have allowed investigations of the ultrastructural organization of the neuropil in three dimensions to uncover the key features of the make-up of synapses in cortical circuits (Denk & Horstmann 2004; Knott et al. 2008; Schalek et al. 2011; Markram et al. 2015; Kasturi et al. 2015; Shapson-Coe et al. 2021; Dorkenwald et al. 2024; Schlegel et al. 2024; Turner et al. 2022; MICrONS Consortium, 2025). There are two major approaches to provide insight brain connectivity, the first is to fully reconstruct a local region of brain using ’dense’ reconstructions (Helmstaedter et al. 2013; Kasthuri et al. 2015; Markram et al. 2015; Lee et al. 2016; Takemura et al. 2017; Zheng et al. 2018; Shapson-Coe et al. 2021; Dorkenwald et al. 2024; Schlegel et al. 2024; MICrONS Consortium, 2025), and the second approach is to undertake ’sparse’ reconstructions of smaller volumes over larger a region of cortex; a method that is better suited to studying the primate cortex (da Costa & Martin 2013). In the current study we have adopted the latter approach to make a detailed quantitative comparison of the synaptic structure in the predominantly input layers of primate primary visual cortex (layer 4C) and in the predominant output layer (layer 3B).

Recently, quantitative studies of synaptic morphology have begun to provide an important component for linking to the physiological and computational outcomes of circuit functioning (Hsu et al. 2017; Bopp et al. 2017; Rodriguez-Moreno et al. 2018; Garcia-Marin et al. 2019; Yakoubi et al. 2019a,b; Ashaber et al. 2020; Matsliah et al. 2024, MICrONS Consortium, 2025). A number of these studies that describe the ultrastructural properties of the cortical columns have been done in rodents (for human see Shapson-Coe et al. 2021) and have followed the dense approach (Kasthuri et al. 2015; Markram et al. 2015; Lee et al. 2016; MICrONS Consortium, 2025). Extrapolating from rodent to primate does not always capture the complexity of primate circuitry because there are a number of major differences between the rodent and primate brains regarding the number, types and distribution of neurons and synapses, the morphological properties of the neurons and their connectivity patterns (DeFelipe et al. 2002; DeFelipe 2011; Barbas 2015; Gilman et al. 2017).

In the current study we have investigated the ultrastructural organization in macaque cortical columns in the visual system. In the visual cortex using a primate model is particularly important, for instance, to understand the properties and organization of the connectivity of neurons in layer 4. The P-pathway makes up 80% of the feedforward (FF) direct thalamic input from the Lateral Geniculate Nucleus (LGN) to the layer 4 primary visual cortex, notably this pathway does not have a counterpart in the rodent. Furthermore, in the achromatic M-pathway of primates, there are functional and modeling constraints that point to the importance of a primate-centric view (Casagrande & Xu, 2004; Callaway, 2005; Horton & Adams, 2005). Recently, we adopted the sparse approach to complement the quantification of the FF thalamocortical (TC) and intracortical (IC) synaptic contributions to different tiers of layer 4C in macaque cortex (Garcia-Marin et al. 2019). Importantly, using this sparse approach we found a substantially stronger TC input to layer 4 in monkey (Garcia-Marin et al. 2019) than previously reports (Peters et al. 1994; Latawiec et al. 2000)

In macaque primary visual cortex (V1) has been proposed that the different inputs, TC vs IC, could have different functional properties, the former being considered ‘drivers’ and the later considered the ‘modulators’ (Sherman, 2005). Previous data has shown that TC synapses have different ultrastructural properties from IC synapses; the TC boutons are larger and establish more synaptic contacts than the IC boutons (McGuire et al. 1984; Freund et al. 1989; Ahmed et al. 1994; Nahami & Erisir 2005; Garcia-Marin et al. 2013). Furthermore, TC boutons, which can be selectively labeled with Vesicular Glutamate Transporter 2 (VGlut2), have been linked to higher probability of neurotransmitter release, a characteristic necessary for the fidelity of synaptic transmission (Covic & Sherman 2011; Fremeau et al. 2004). In contrast, the IC boutons, which can be identified with Vesicular Glutamate Transporter 1 (VGlut1), are smaller and have been linked to low probability of transmitter release and various types of plasticity, especially potentiation (Balschun et al. 2010; Covic & Sherman 2011; Fremeau et al. 2004). The goal of the current study was undertake a detailed ultrastructural analysis of the synapses in two different layers of macaque V1, layer 4C, the main TC recipient layer, with different physiological properties in each sublayer (Tootell et al. 1988, Hawken & Parker 1984; Hawken et al. 1988; Saul et al. 2005), and compare them with layer 3B, a principal output layer and with very sparse TC input (Garcia-Marin et al. 2013). We propose that the unique ultrastructural properties of the synapses in the different layers would provide a signature of circuit processing, and this also would provide the initial foundation for components of realistic functional models of the circuits in the input layers and the supragranular cortical layers. Both the different and the common characteristic in both layers could give us insight about the circuit connectivity of cortex.

## MATERIALS AND METHODS

### Macaque brain tissue

Three macaque monkeys (one male *M. fascicularis*, age 3.6 years and two female *M. nemestrina*, ages 7.7 and 18.5 years), previously used for anesthetized electrophysiological recordings, were used in this study. Animals were prepared for recording as described elsewhere (Xing et al. 2004; Solomon et al. 2004). After 4–5 days of recordings, experiments were terminated by intravenous injection of a lethal dose of pentobarbital (60 mg/kg), and brain death was determined by a flat electroencephalogram. Subsequently, animals were transcardially perfused with heparinized 0.01 M phosphate-buffered saline (PBS; pH 7.4) followed by 4% PFA + 0.125% Glutaraldehyde (Glu) in 0.1 M PB. Some blocks of V1 were removed for track reconstruction of the recording locations of electrophysiologically characterized neurons, and the remaining V1 tissue was cut into small blocks and postfixed in the same fixative for 24–72 hours at 4°C. After fixation, coronal sections of 100 and 50 µm from the opercular region of V1 – representing eccentricities of 2–5 degrees – were prepared using a vibratome. The remaining sections and blocks were immersed in graded sucrose solutions, and were stored in a cryoprotectant solution at -20°C.

The layers of area V1 were identified according to Brodmann’s (Brodmann 1909) nomenclature, modified by Lund (Lund 1973). This system distinguishes four subdivisions of layer 4, namely, 4A, 4B, 4Cα, and 4Cβ. While Brodmann’s (1909) laminar framework for V1 is commonly employed, it should be noted that the subdivision of layer 4 into three sublayers (4A, 4B, 4C) does not accurately correspond to the fact that only layer 4C aligns with layer 4 in other regions, as proposed by Hassler in 1967. In this context, Brodmann’s layers 4A and 4B are regarded as sublayers of layer 3. Despite the adoption of Hassler’s terminology in some contemporary studies of V1 in primates (see Balaram et al. 2014; Balaram & Kaas 2014), there are still influential investigations, particularly those delving into circuitry and function (see Adams et al. 2007; Sincich et al. 2003; Federer et al. 2013; Callaway 1998), that adhere to Brodmann’s nomenclature. To facilitate comparison with other V1 studies in both human and non-human primates utilizing varied nomenclature, our sampling regions included layer 3B, layer 4Cα (4A), and layer 4Cβ (4B), maintaining Brodmann’s nomenclature throughout the text.

All experimental procedures were approved by the New York University Institutional Animal Use and Care Committee and were conducted in strict compliance with the National Institutes of Health (NIH) guidelines for the care and experimental use of animals in research.

#### Tissue processing

Free-floating sections were treated in 1% osmium tetroxide in 0.1 M PB for 45 minutes and rinsed in 0.1 m PB. The following steps were undertaken in a Pelco BioWave (Ted Pella, Inc., Redding, CA) that shortens by up to 95% the traditional bench processing time while preserving and improving the preservation of the membranes (Webster, 2014). Dehydration was initiated with a series of 30% and 50% ethanol, followed by 3 series of uranyl acetate 2% in 70% ethanol, followed by 95% ethanol, twice in 100% ethanol and twice in acetone (each step 40 seconds). The infiltration with 1:1 acetone:araldite took 3 min, and finally the polymerization took place overnight at room temperature with 100% araldite. The following day the sections were flat embedded in Araldite resin, the cured for 48 h at 60 deg C.

A correlative light and electron microscopy method (DeFelipe & Fairen 1993) was used to identify the regions of interests (layers 3B, 4Cα and 4Cβ) (Fig. 1A). Sections that included all the layers in V1 were photographed under the light microscope, glued on a pre-polymerized resin blocks in a beam capsule (Fig. 1A) and then cut into serial semithin (1 µm thick, Fig. 1B) sections with an Ultracut (RMC, Inc. MT6000-XL, Wien, Austria) ultramicrotome. The semithin sections were stained with 1% toluidine blue in 1% borax, and examined under the light microscope, to identify the laminar boundaries based on neuronal density (Fig. 1B-C). Once the regions of interests were identified in the semithin sections (Fig. 1C), the resin block, was trimmed to a smaller block that included layers 1-2-3 or 4Cα and 4Cβ (Fig. 1D-F). For the layer 3B regions we sampled at 300-400 µm from the pial surface. From the layer 4C block, we sampled from the upper and lower parts of layer 4C, for layer 4Cα and 4Cβ, respectively, with a minimum of 100 µm between these two regions (Fig. 1D-F).

**Figure 1:**
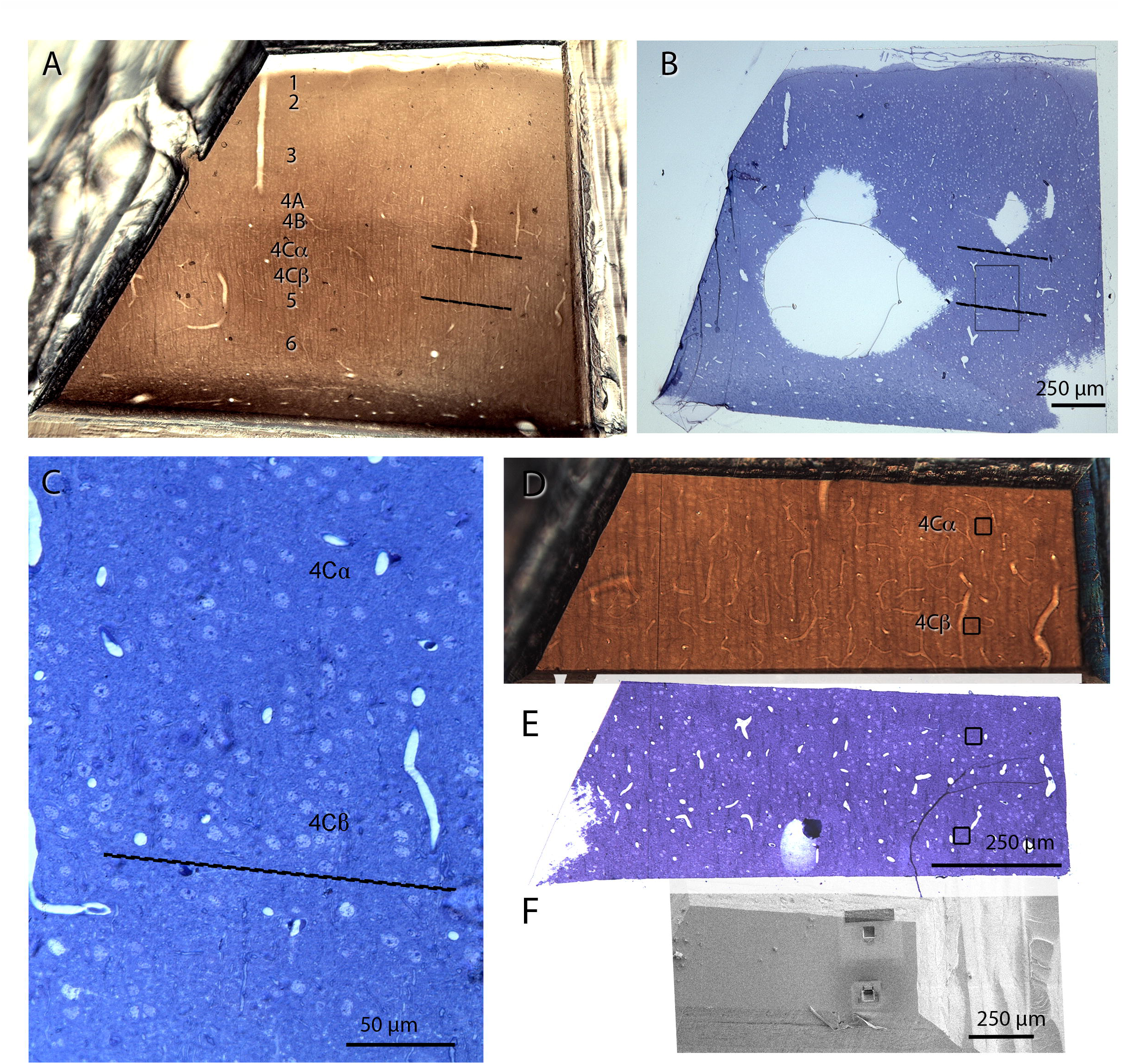
Correlative light and electron microscopy to determine the location of layers 3B, 4Cα, and 4Cβ. A) In the thick section (100 µm) fixed with osmium, a darker band can be observed at the level of layer 4. Approximate boundaries of layer 4Cα and 4Cβ indicated with black lines B) Semithin section (1 µm) from the surface of the block stained with toluidine blue, aligned with the thick section to find the same laminar boundaries. C) Higher magnification of the box in image Bthat shows high neuronal density allowing the unequivocal identification of layers 4Cα and 4Cβ. D) A thick section fixed with osmium trimmed to include layer 4C. E) The last semithin section (which corresponds to the section immediately adjacent to the block surface) was examined under light microscope and photographed to accurately locate the region to be examined. F) Image taken with the SEM from the surface of the block containing the layers to be analyzed (4Cα and 4Cβ).

The degree of tissue shrinkage was measured by comparing sections in x, y, and *z* dimensions before and after tissue preparation with the aid of Olympus VS120. The average linear shrinkage (S) was calculated by using the following formula: S = (A-0)/A, where A is the absolute value before processing and 0 is the observed value after processing. This yielded an average linear shrinkage of 3%, 3% and 9.5%, in x, y, and z, respectively, for EM.

### Focused Ion Beam/ Scanning Electron Microscope image acquisition

The three-dimensional images were obtained using the Helios Nanolab DualBeam (FIB/SEM) from FEI (Hillsboro, OR, USA) at the Simons Electron Microscopy Center located at the New York Structural Biology Center. This microscope utilizes unique DualBeam technology, which combines Focused Ion Beam (FIB) milling and Scanning Electron Microscope (SEM) imaging. The block face is imaged with a scanning electron beam and then the surface is milled; the sequential process of imaging and milling provides serial thin sections of tissue, thereby allowing the visualization and reconstruction of the 3D spatial organization of elements at the ultrastructural level. For the current study we used a milling thickness of 20 nm, while imaging of the exposed surface was obtained at 1.8 kV acceleration potential using the in-column energy selective backscattered (EsB) electron detector. The aperture size was 30 µm, and the retarding potential of the EsB grid was 1500 V. The imaging and milling processes were repeated in a fully automated way to obtain long series of images (a stack) that represented a 3D sample of the tissue. Each stack had x,y dimensions of 4096 x 3536 pixels, with an x,y resolution ranging from 2.93 to 3.38 nm/pix, and z dimensions ranging from 223 to 499 images, with a z step size of 20 nm (Fig. 2A).

**Figure 2:**
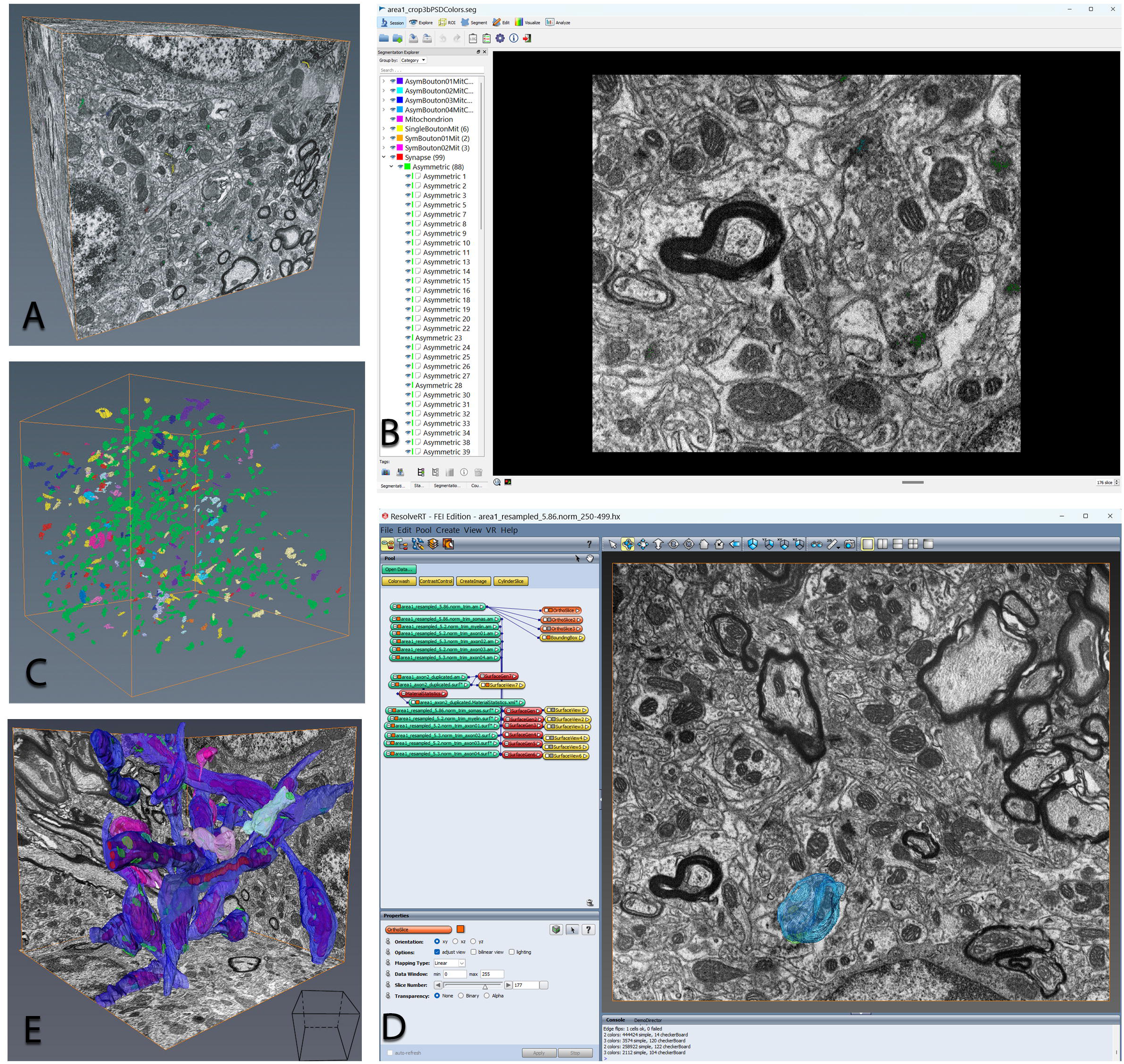
Three dimensional reconstruction and analysis of synaptic elements in each block of Vl cortical tissue. A) Block of macaque Vl from layer 4Cβ obtained from projecting 500 FIBSEM images. Each image resolution was 3.7 nm/pix in x,y and 20 nm in z. Total volume 944 µm^3^• B) Screenshot of the ESPina software interface that allows the identification and segmentation of the synapses. C) 3D reconstructed synaptic junctions, each displayed by different colors representing different axonal categories: green, asymmetric synapses (SSBm-); red, symmetric synapses; yellow, asymmetric synapses with mitochondria (SSBm+); other colors, asymmetric synapses belonging to an axon that established multiple synapses (MSBm+); D) Screenshot of the AMIRA software with same stack of images, where the axons were reconstructed in 3D. An example of one of these reconstructed axons in colored in blue. E) AMIRA 3D reconstructions of the stack in A, the different axon categories were assigned to different colors for the purpose of representation: MSBm+, blue; MSBm-, cyan; SSBm+, dark pink; SSBm-, light pink; mitochondria (red) are observed by transparency inside the axons, and the synapses are colored in green.

### Measuring synaptic density using ESPina

To quantify the total density of synapses in the neuropil of layers 3B, 4Cα and, 4Cβ the original stacks were cropped into non-overlapping subregions for analysis and large blood vessels and neurons were excluded. We used 18, 17 and 18 image stack subregions from three monkeys, with a mean total volume sampled of 126 µm^3^ (38 - 192 µm^3^), 104 µm^3^ (20 - 186 µm^3^) and 95 µm^3^ (35 - 200 µm^3^) in layers 3B, 4Cα and 4Cβ, respectively (Suppl. Table 1, 2, 3 for layers 3B, 4Cα and 4Cβ, respectively).

Identification of synapses in 3D allowed a complete classification of all synapses, regardless of the plane of section. A major advantage of the 3D technique is that it provides a classification of all the synapses while the 2D TEM approach leaves approximately 40-60% of synapses uncharacterized (DeFelipe et al. 1999; Kubota et al. 2009; Garcia-Marin et al. 2019). Synapses with a prominent PSD (Post Synaptic Density) were classified as asymmetric (Asym), and synapses with a thin PSD were classified as symmetric (Sym) (Colonnier 1968; for review see Colonnier 1981; Peters et al. 1991; Peters & Palay 1996).

With the aid of ESPina software (Espina Interactive Neuron Analyzer, 2.4.1; Madrid, Spain; https://cajalbbp.es/espina/), the observer can navigate through the stack of images and unambiguously identify every single synapse as asymmetric of symmetric based on their PSDs (Fig. 2B). The manually identified synapses were automatically segmented and reconstructed in 3D by ESPina based on the gray levels and connectivity (Morales et al. 2011) (Fig. 2B-C). The expert observer then confirmed this segmentation. To quantify the number of synapses per volume, a 3D counting frame was defined within the stack in ESPina. This unbiased “counting brick” was a rectangular prism bounded by three acceptance planes and three exclusion planes (Howard & Reed 2005; Morales et al. 2011). All objects within the counting brick or intersecting any of the acceptance planes (defined as the front, top, and left sides of the stack subregion) were counted, while any object outside the counting brick or intersecting any of the exclusion planes (back, right, and bottom sides) were not included in the quantification.

After segmenting all the synapses, they were further assigned to different boutons categories: MultiSynaptic Boutons (MSBs), those boutons that make synaptic contacts with two or more postsynaptic structures (Peters et al. 1991; Jones et al. 1997) and single-synapse boutons (SSBs), those that establish a single synapse with a postsynaptic element (Fig. 2C). MSBs have been observed in different species (ferrets, cats, mice, rats, monkeys) and areas (visual, somatosensory, motor, frontal cortices) (see Winfield & Powell 1983; Nahmani & Erisir 2005; Graziano et al. 2008; Bopp et al. 2017; Hsu et al, 2017, Garcia-Marin et al. 2019). Further, each category was subdivided by presence or absence of mitochondria (m+ and m-, respectively). Mitochondria are found in the presynaptic axon terminal of neurons, playing a crucial role in providing the substantial energy required for the processes involved in neurotransmission, where the terminal is responsible for releasing neurotransmitters (review in Duarte et al. 2023).

In addition, the postsynaptic element of each synapse was identified by the characteristics in 3D. We distinguished dendritic shaft by the presence of microtubules, mitochondria, and smooth endoplasmic reticulum in the cytoplasm, and dendritic spines. Spines are small protrusions from a dendrite, that contains a cytoplasm with floccular material sometimes occupied by a specialized form of endoplasmic reticulum, the spine apparatus (Peters et al. 1991). Finally, synaptic junction shapes were also examined with ESPina and were classified into four morphological categories: macular, horseshoe-shaped, or perforated (Montero-Crespo et al. 2021). The boutons establishing symmetric synapses were assigned to a single category. More detailed analysis of the symmetric synapses will be included in a future study.

### 3D reconstruction of boutons using AMIRA

Using the images from the FIB/SEM, we identified a total of 159 boutons that were fully included in the stacks and reconstructed their profiles in 3D using Amira 3D software (Fig. 2D) (Thermo Fisher Scientific – FEI). To calculate the volume occupied by the different compartments we used separate “materials” to identify axonal membrane, mitochondria, and PSDs. We traced the profile of each object in each image plane across the full extent of each bouton and generated a 3D reconstruction for each material (Fig 2E, 3A, Suppl Fig 1). To obtain an accurate estimate of the bouton volume, the axonal fiber adjacent to the bouton was cropped from the 3D reconstruction (Fig. 3B-C, black lines). Axonal boutons were classified following the same classification as described above in measurements of synaptic density using ESPina: SSBm+ (single-synaptic bouton with mitochondria, Fig. 3C), MSBm+ (multisynaptic bouton with mitochondria, Fig. 3B), SSBm-(single-synaptic bouton without mitochondria, Fig. 3E), and MSBm-(multisynaptic bouton without mitochondria, Fig 3D). The following parameters were calculated for each category: bouton volume, mitochondria volume, proportion of bouton volume occupied by mitochondria, synapses per bouton and PSD area. The PSD area was calculated by dividing the PSD volume obtained from the 3D reconstruction (using AMIRA) by an average PSD thickness. The average thickness was calculated with ImageJ (NIH) by taking measurements of PSDs that were oriented with their faces orthogonal to the xy-plane. Twenty-five orthogonal PSDs were identified and measured. With each PSD, 3 thick measurements – two near the ends of the PSD and one in the center – were taken for every image plane that contained the PSD structure (no significant differences were found between the two edges and center, (ANOVA, p 0.256). The mean value of the PSD thickness was 30.5 ± 4.7 nm (mean ± 1 SD, n = 75) (Suppl Fig. 2).

**Figure 3:**
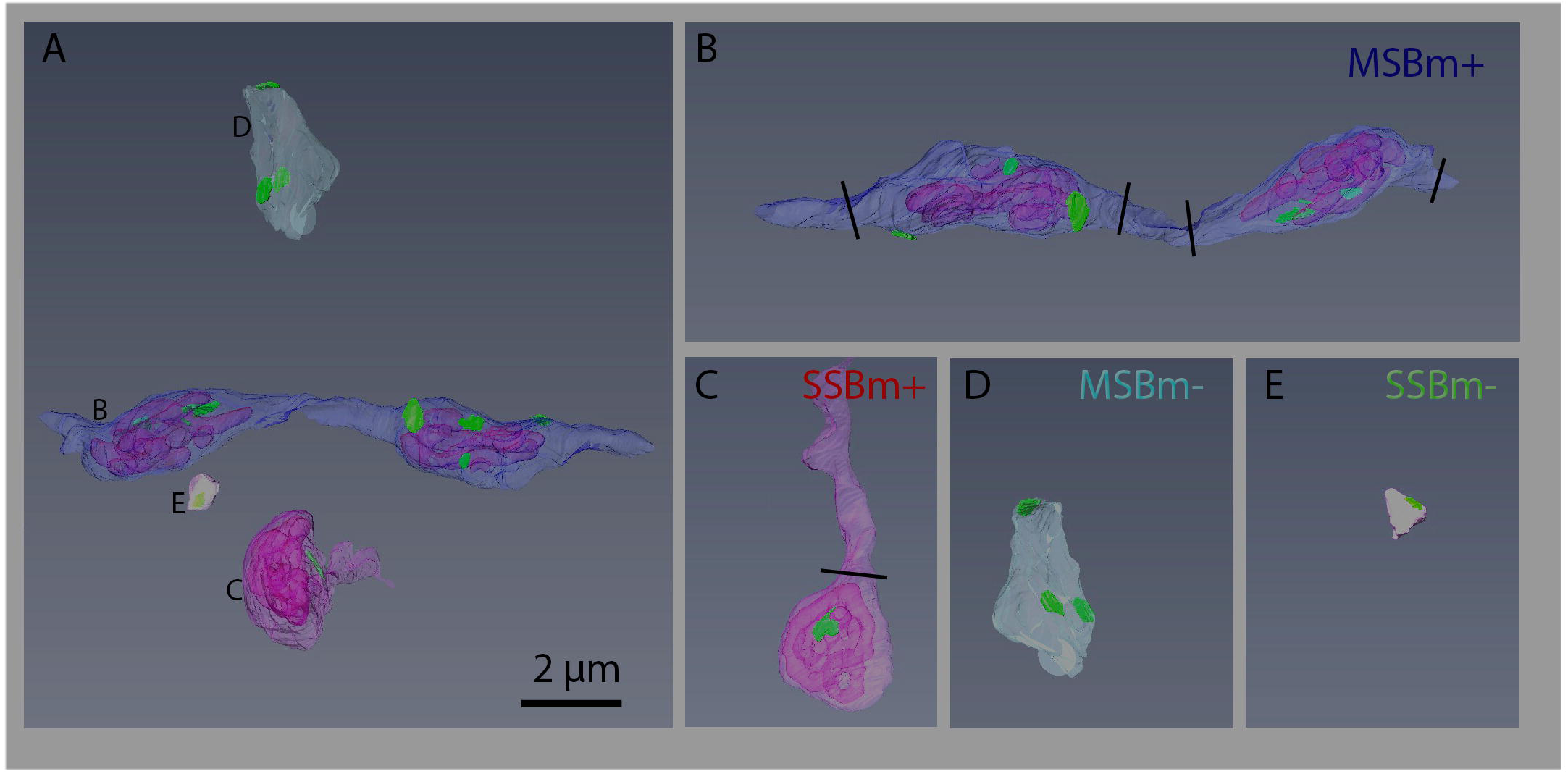
Individual boutons categories and the arrangements of their components. A) Examples of different axonal 3D morphologies reconstructed with AMIRA. B) MSBm+, category that incorporates boutons with mitochondria establishing more than 1 asymmetric synapses. C) SSBm+, category that includes boutons with mitochondria establishing 1 asymmetric synapse. D) MSBm-, category to classify boutons without mitochondria establishing more than 1 asymmetric synapses. E) SSBm-, category that includes boutons without mitochondria establishing 1 asymmetric synapse. Synapses in green, mitochondria in transparent pink inside the boutons. Black lines indicate cutting regions to calculate the volume only in the terminal area.

#### Estimation of the volume fraction for cortical components

Three semithin sections, each 1 μm thick and stained with 1% toluidine blue, were selected from every case. These sections were utilized to assess the volume fraction of blood vessels, cell bodies, and neuropil within each layer. The estimation process followed the Cavalieri principle (Gundersen et al., 1988), employing point counting through the integrated Stereo Investigator stereological package (Version 8.0, MicroBrightField Inc, VT, USA) connected to an Olympus light microscope (Olympus, Bellerup, Denmark) at a magnification of 40x. Over each semithin section, a grid with points associated with an area of 900 µm^2^ was overlaid to determine the volume fraction (Vv) occupied by the principal tissue elements: blood vessels, glia, neurons, and neuropil. The volume fraction occupied by neuropil was calculated using the formula: Vv neuropil = 100 – (Vv blood vessels + Vv cells (glia + neurons)) (Fig 4A-B).

**Figure 4:**
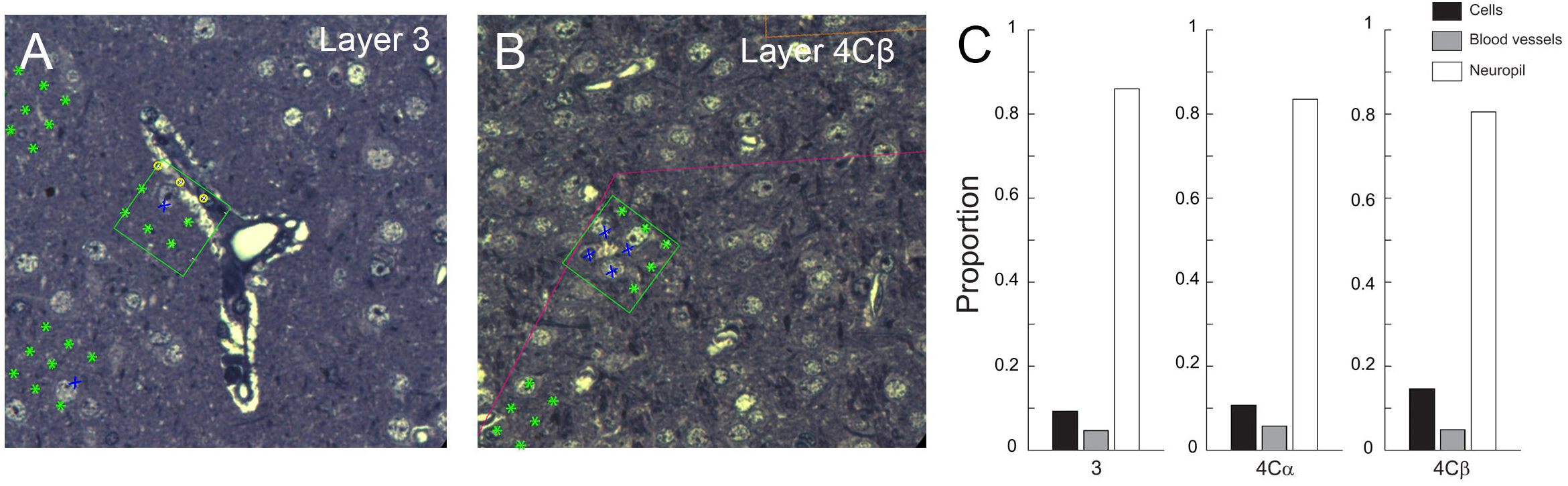
Quantification of the different elements of the neuropil of layers 3B, 4Cα, 4Cβ. A-B) Semithin sections of layers 3B and 4Cα with the counting grids superimposed to stereologically estimate the volume fraction occupied by the different elements: neuropil (green asterisks), cells (neurons and glial bodies, blue crosses), and blood vessels (yellow circles). C) Graph showing the Vv occupied by cells (neuronal and glial somata), blood vessels, and neuropil in 3, 4Cα, and 4Cβ of macaque Vl. The Vv by the neuropil was significantly different in the three layers **(KW,** p==0.0002). Layer 4CP had significantly less neuropil than layer 3 and layer 4Cα (post-hoc: 4Cβ vs 4Cα, ***; 4CP vs 3, ***, 3 vs 4Cα, ns). The Vv of cells was also significantly different by layers (KW, p<0.0001; layer 4CP had more cells than layer 3 and 4Cα (post-hoc: 4Cα vs 4Cβ, ***; 4CP vs 3, ***). No significant difference was found in the Vv of blood vessels in the different layers (KW, p==0.1547, ns). **** p < 0.0001; *** p < 0.001; ** p < 0.01; * p < 0.05; ns p > 0.05

### Statistical analysis

Statistical comparisons of synaptic density, morphological parameters, mitochondrial content, and the volume fraction (Vv) occupied by cortical elements were performed using either parametric or non-parametric tests, depending on data distribution. When data were normally distributed and passed the test for homogeneity of variances (Bartlett’s test), parametric tests (t-test or one-way ANOVA) were used, followed by Tukey’s Honestly Significant Difference (HSD) test for post hoc pairwise comparisons. For non-normally distributed variables, non-parametric tests (Mann–Whitney (MW) U test or Kruskal– Wallis (KW) test) were applied, followed by Dunn’s test with Sidak correction for post hoc pairwise comparisons. To analyze bouton volume categories and PSD areas across layers, we plotted cumulative frequency distributions and used the Kolmogorov–Smirnov (K–S) test to assess whether the distributions differed significantly. For categorical data, contingency tables were analysed using chi-square (χ²) tests. When multiple pairwise comparisons were required, we applied a Bonferroni correction by dividing the significance level by the number of comparisons. Statistical significance was defined as *p* < 0.05. Exact *p*-values were reported for all the statistical tests. Data were presented as mean ± standard error of the mean (SEM) or standard deviation (SD), as appropriate. All statistical analyses were performed using MATLAB R2022a (MathWorks, Natick, MA) and GraphPad Prism 5.0 (GraphPad Software, Boston, MA).

## RESULTS

Automatic serial EM image acquisition with the FIB/SEM allowed us to study a large volume of brain tissue at different locations, to address the question of whether different layers have a distinct neuropil composition. We classified 1430, 939, and 1537 synapses in layers 3B, 4Cα and 4Cβ, respectively. Further, we fully reconstructed in 3D, 45, 48 and 63 boutons in layers 3B, 4Cα and 4Cβ, respectively.

### Volume fraction of cortical elements

Across the three layers, the average values of the estimated Vv occupied by cells (neurons and glial cell bodies), blood vessels, and neuropil were 12%, 5%, and 83%, respectively. When the values were compared by layers, we found that the Vv by the neuropil was significantly different in the three layers (KW, p<0.0002); layer 4Cβ had significantly less neuropil (80.50%±0.0075) than layer 4Cα (83.52%±0.0012), and layer 3B (85.99%±0.0028) (4Cβ vs 4Cα, p =0.0391; 4Cβ vs 3, p=0.001, 3 vs 4Cα, p=0.1043, ns) (Figure 4). The Vv of cells was also significantly different across layers (KW, p<0.0001), layer 4Cβ had significantly more cells (14.61%±0.0097) than 4Cα (10.74%±0.0099) and layer3 (9.3%±0.0038) and (4Cα vs 4Cβ, p=0.0063; 4Cβ vs 3, p=0.0002; 3 vs 4Cα, ns) (Figure 4, Table 1). No significant differences were found in the Vv of blood vessels in the different layers (KW, p=0.1547) (4.7%±0.0031, 5.75%±0.0046, 4.89%±0.0040, for layer 3, 4Cα and 4Cβ, respectively).

**Table 1:** Synaptic density (number synapses/µm^3^) in layer 3B, 4Cα, and 4Cβ (asymmetric (asym), symmetric (sym), and total (asym+sym)). Proportion of symmetric and asymmetric. Two (for layer 3B) and three for layers 4cα and 4cβ animals contributed to the mean values shown in this table. There were 18, 17 and 18 individual counting frames contributing to the values in this table. CF: counting frame.

| | 3B | 4Cα | 4Cβ |
| --- | --- | --- | --- |
| No. asym<br>(included in CFs) | 1202 | 799 | 1333 |
| No. sym<br>(included in all CFs) | 228 | 140 | 204 |
| No. total<br>(included in CFs) | 1430 | 939 | 1537 |
| % asym | 84% | 85% | 87% |
| % sym | 16% | 15% | 13% |
| Mean CF volume ($\mu\text{m}^3$) | 126 | 104 | 95 |
| No. Asym /$\mu\text{m}^3$<br>(mean±SEM) | 0.507±0.022 | 0.404±0.021 | 0.653±0.032 |
| No. Sym /$\mu\text{m}^3$<br>(mean±SEM) | 0.082±0.004 | 0.071±0.004 | 0.098±0.005 |
| No. total synapses/$\mu\text{m}^3$<br>(mean±SEM) | 0.589±0.026 | 0.476±0.024 | 0.751±0.037 |

### Quantification of the Synaptic Density in different layers

Stacks of images obtained using the FIB/SEM were analyzed using ESPina software, which provides a segmentation of synapses in 3D (Morales et al. 2011) hence allowing for a more accurate estimation of the number, size, and arrangement of synapses than 2D segmentation (DeFelipe et al. 1999; Kubota et al. 2009; Garcia-Marin et al. 2019). Recently, using this methodology, we have reported that the synaptic density in 4Cβ was 1.5 times greater than the synaptic density in 4Cα (0.75 syn/μm^3^ vs 0.476 syn/μm^3^ (Table 1, Supplementary Table 2-3) (Garcia-Marin et al., 2019). In the current study we expanded this work to compare the synaptic density of the main TC recipient layers (4Cα and 4Cβ), with layer 3B, a non-TC recipient layer (except for the CO-patches) (Table 1, Supplementary Table 1). We found that the total synaptic density in layer 3B was 0.59 syn/μm^3^ ± 0.026– with 1.3 times fewer synapses than in 4Cβ (Fig 5A) and 1.2 more synapses than 4Cα (Table 2, Figure 5A, Anova, p<0.0001, post-hoc comparisons: L3B vs 4Cα (p=0.0464), L3B vs 4Cβ (p=0.0016), and 4Cα vs 4Cβ (p<0.0001).

**Figure 5:**
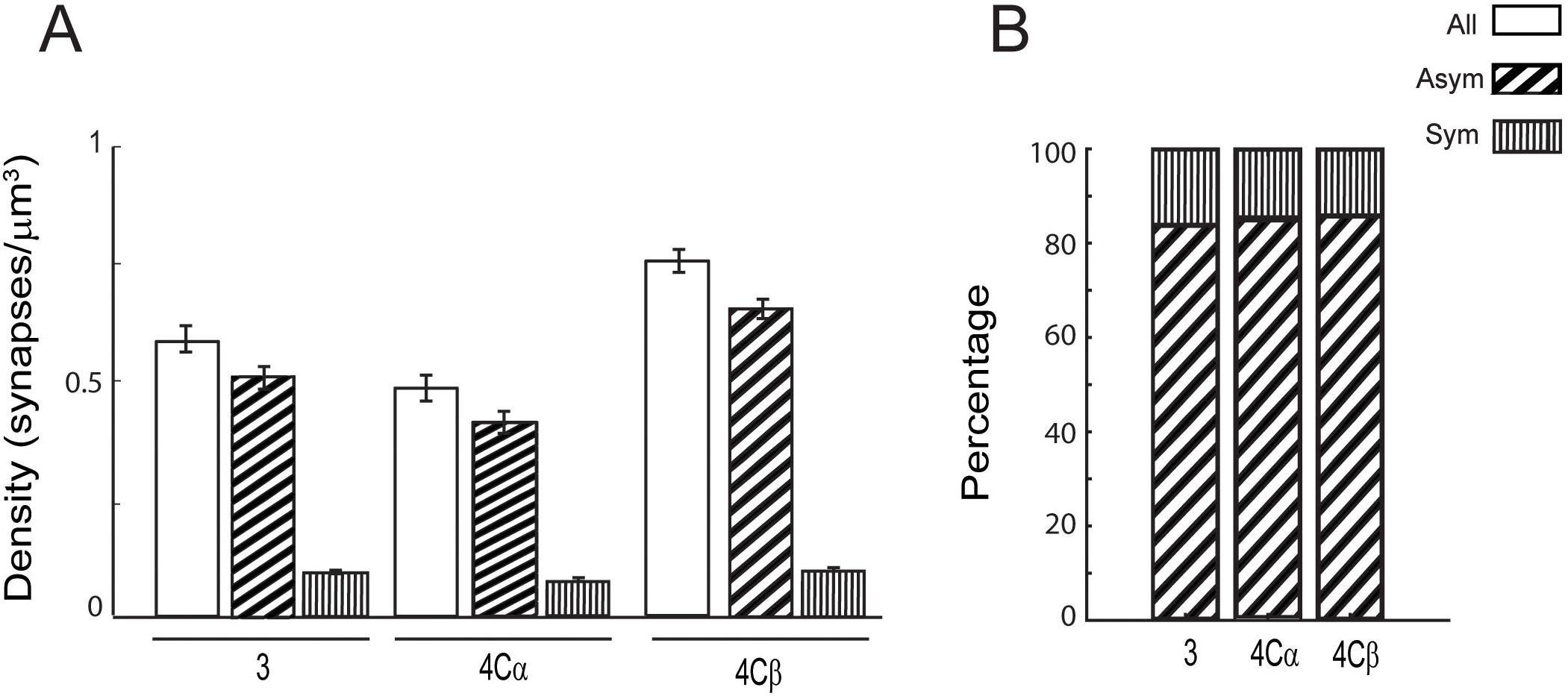
Synaptic density in different cortical layers. A) Bar graph showing the synaptic density for asymmetric and symmetric synapses and the sum of both (all) (mean± SD) in layer 3, 4Cα, and 4Cβ of macaque Vl. The statistical analyses showed that there were differences in the total synaptic density by layers (Anova, p<0.0001), post-hoc: L3B vs 4Cα (*), L3B vs 4Cβ (**), and 4Cα vs 4Cβ (****). The density of asymmetric synapses also significantly varied by layers (Anova, p<0.0001, post-hoc: L3B vs 4Cα (*), L3B vs 4Cβ (***), and 4Cα vs 4Cβ (****). The density of symmetric synapses was also significantly different by layers 4Cα vs 4Cβ (Anova, p<0.0001, post-hoc: L3B vs 4Cα (ns), L3B vs 4Cβ (*), and 4Cα vs 4Cβ (***). B) The proportion of symmetric and asymmetric synapses in layers 3, 4Cα, and 4Cβ of macaque Vl were not significantly different (x^2^, ns). **** p < 0.0001; *** p < 0.001; ** p < 0.01; * p < 0.05; ns p > 0.05

**Table 2:** Classification of the asymmetric synapses from ESPina in the different bouton categories based on the number of synapses that they established and the presence of mitochondria: SSBm-(a bouton making one single asymmetric contact without mitochondria), SSBm+ (a bouton establishing one single asymmetric contact with mitochondria), MSBm+ (a bouton making ≥2 asymmetric contacts with mitochondria), and MSBm-(a bouton making ≥2 asymmetric contacts without mitochondria. Each entry in the category rows gives the total number and the percent of total in parentheses. The last row (Combined) shows the data combined across all four categories.

| Bouton Category | All layers | 3B | 4Cα | 4Cβ |
| --- | --- | --- | --- | --- |
| MSBm+ | 276<br>(8.5%) | 71<br>(6.4%) | 34<br>(4.3%) | 171<br>(12.8%) |
| SSBm+ | 709<br>(21.9%) | 287<br>(25.9%) | 239<br>(29.9%) | 183<br>(13.7%) |
| SSBm- | 2225<br>(68.7%) | 750<br>(67.7%) | 521<br>(65.1%) | 954<br>(71.6%) |
| MSBm- | 31<br>(0.9 %) | 0<br>(0%) | 6<br>(0.7%) | 25<br>(1.9%) |
| **Combined** | **3241** | **1108** | **800** | **1333** |

( $\chi^2$ , \*\*\*\*, MSBm+: Bonferroni comparison: 3 vs 4Cα (\*\*\*\*), vs 4Cβ (\*\*\*\*, 4Cα vs 4Cβ (\*\*\*\*); SSBm+: Bonferroni comparison: 3 vs 4Cα (\*\*), vs 4Cβ (\*\*\*\*), 4Cα vs 4Cβ (\*\*\*); SSBm-: Bonferroni comparison: 3 vs 4Cα (\*\*\*\*), vs 4Cβ (\*\*\*\*), 4Cα vs 4Cβ (\*\*\*\*); \*\*\*\* p < 0.0001; \*\*\* p < 0.001; \*\* p < 0.01; \* p < 0.05; ns p > 0.05

The 3D segmentation allowed unambiguous classification of the synapses into asymmetric (with a thick PSD) or symmetric (thin PSD). The density of asymmetric (As) significantly varied by layers, 0.507 syn/μm^3^ ± 0.022, 0.404 syn/μm^3^ ± 0.021, and 0.653 syn/μm^3^ ± 0.032 syn/μm^3^ for layer 3B, 4Cα and, 4Cβ, respectively (Table 1, Figure 5A, Anova, p<0.0001, post-hoc: L3B vs 4Cα (p=0.0357), L3B vs 4Cβ (p=0.0009), and 4Cα vs 4Cβ (p<0.0001)). The density of symmetric (Sy) also varied by layers, 0.082 syn/μm^3^ ± 0.004, 0.071 syn/μm^3^ ± 0.004, and 0.098 syn/μm^3^ ± 0.005 for layer 3B, 4Cα and, 4Cβ, (Table1, Fig 5A, Anova, p<0.0001, post-hoc: L3B vs 4Cα (p=ns), L3B vs 4Cβ (p=0.0395), and 4Cα vs 4Cβ (p=0.0002)). When the proportion of asymmetric and symmetric synapses was calculated, the ratios were very similar across layers 86:14% in layer 3B, 85:15% in layer 4Cα, and 87:13% in layer 4Cβ (Table 1, Fig. 5B). (Table 1, Fig 5B, (χ², p=0.117, ns). These results suggest that the As:Sy synapse ratio is overall consistent across layers despite variations in absolute synapse density.

#### Diversity and distribution of asymmetric boutons

As the proportion of asymmetric versus symmetric synapses was constant across the layers, the next step was to investigate whether the proportion of different axonal bouton categories was also constant across layers. Each asymmetric synapse was assigned to a axonal bouton category based on the number of synapses that it established and by the presence or absence of mitochondria: SSBm-(a bouton making one single asymmetric contact without mitochondria) (Fig. 3E), SSBm+ (a bouton establishing one single asymmetric contact with mitochondria) (Fig. 3C), MSBm+ (a bouton making ≥2 asymmetric contacts with mitochondria) (Fig. 3B), and MSBm-(a bouton making ≥2 asymmetric contacts without mitochondria) (Fig. 3D).

There were clear differences in the morphology of the 4 bouton categories (Fig. 3). The SSBm+, SSBm-, and MSBm-boutons were bulbs of different sizes at the end of the axon that contains vesicles. These boutons are known as ‘terminal boutons’ (Peters et al., 1991) (Fig 3C-E). The MSBm+ boutons were swellings found along the length of the fiber; these boutons are known as ‘*en passant’* boutons (Peters et al., 1991) (Fig. 3B, Fig. 12A-H). To estimate the spatial characteristics of the ‘*en passant’* boutons we calculated the interbouton distance from 6 reconstructed fibers that established 11 MSBm+ boutons. The average interbouton distance was 2.3±1.2 µm (mean ±1SD; range 0.99–3.98), measured as the distance between the end of a varicosity and the beginning of the next varicosity.

The three axonal bouton categories – MSBm+, SSBm+, SSBm-– were found in all the layers (Fig. 6A, Table 2) while MSBm-synapses were found in layers 4Cα, 4Cβ but not in layer 3B. When considering the three layers together, the most abundant bouton was the SSBm-category (69%), followed by SSBm+(22%), MSBm+ (9%), while MSBm-synapses accounted for <1% (Table 2). This overall distribution was observed in layer 3B and 4Cα, and 4Cβ, but with different proportions.

**Figure 6:**
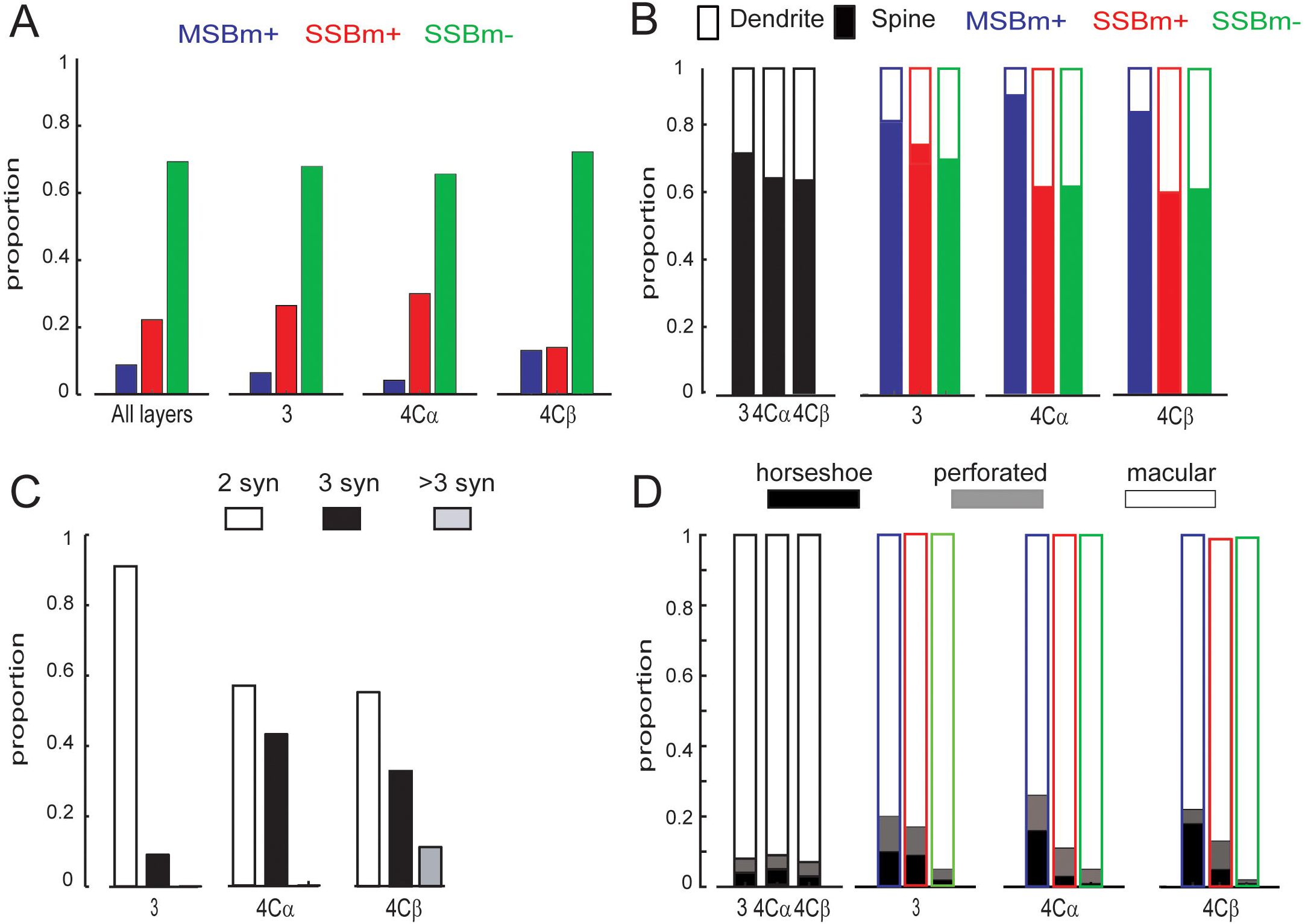
Distribution of synaptic boutons categories across cortical layers and structure of the postsynaptic densities. A) Proportion of the different types of asymmetric boutons: MS8m+ {blue), SSBm+ (red), and SSBm- (green) in all layers combined, layer 3, 4Cα, and 4Cβ (x^2^, p<0.0001. For SSBm-: 3 vs 4Cα (****), vs 4Cβ (****), 4Cα vs 4Cβ (****). For SSBm+, post-hoc: 3 vs 4Cα (***), vs 4Cβ (****), 4Cα vs 4Cβ (***)). For MS8m+, post-hoc: 3 vs 4Cα (***), vs 4Cβ (****), 4Cα vs 4Cβ (****). 8) Proportion of the preferred postsynaptic targets: spine (filled) or dendrite (empty) of each bouton category in all layers combined, (x^2^, p<0.0001, post-hoc: 3 vs 4Cα (****),vs 4Cβ (****), 4Cα vs 4Cβ (****);inlayer 3: (x^2^, p<0.0001, post-hoc: 3 vs 4Cα (****), vs 4Cβ (****), 4Cα vs 4Cβ (****);inlayer 4Cα: (x^2^, ****, post-hoc: 3 vs 4Cα (****), vs 4Cβ (****), 4Cα vs 4Cβ (p<0.0001); in layer 4Cβ: (x^2^, p<0.0001, post-hoc: 3 vs 4Cα (*), vs 4Cβ (****), 4Cα vs 4Cβ (****) C) Proportion of MS8m+ boutons establishing two or more synapses in each layer (x^2^, **; for 2 synapses: 3 vs 4Cα (****), 38 vs 4Cβ (p = 0.6016, ns); 4Cα vs 4Cβ (****); For 3 synapses: 38 vs 4Cα (ns), 38 and 4Cβ (****), 4Cα and 4Cβ (***). D) Proportion of different PSD morphologies (horseshoe, perforated, and macular) in the different axonal categories in all layers combined (x^2^, p=0.089, ns). In layer 38, x^2^, p=0.0001; macular: post-hoc: MS8m+ vs SSBm+ (****), vs SSBm- (**), SSBm+ vs SSBm- (****); horseshoe: post-hoc: MS8m+ vs SSBm+ (*), vs SSBm- (*), SSBm+ vs SSBm- (ns)); perforated, post-hoc: MS8m+ vs SSBm+ (ns), vs SSBm(*), SSBm+ vs SSBm- (ns)). In layer 4Cα (x^2^, ****; macular, post-hoc: MS8m+ vs SSBm+ (****), vs SSBm- (****), SSBm+ vs SSBm- (****)). Perforated: post-hoc: MS8m+ vs SSBm+ (ns), vs SSBm- (*), SSBm+ vs SSBm- (ns)). Horseshoe, post-hoc: 3 vs 4Cα (ns), vs 4Cβ (ns), 4Cα vs 4Cβ (, ns)). In layer 4Cβ (x^2^, p=0.0001; mact1lar, post-hoc: MSBm+ vs SSBm+ (ns), vs SSBm- (****), SSBn1+ vs SSBm(****). Horseshoe, post-hoc: MSB+ bo11tons MSBm+ vs SSBm+ (***), vs SSBm- (*), SSBm+ vs SSBm- (ns). Perforated, post-hoc: MSBm+ vs SSBm+ (ns), vs SSBm- (ns), SSBm+ vs SSBm- (ns). **** p < 0.0001; *** p < 0.001; ** p < 0.01; * p < 0.05; ns p > 0.05

The most abundant class of synapses, SSBm-, accounted for between 65% to 72% of the total number of synapses in the three layers, nonetheless there were small but significant laminar differences with layer 4Cβ having more SSBm- (72%) than in layer 3B (68%) and 4Cα (65.1 %), (Fig. 6A, Table 2, χ², p<0.0001, Bonferroni comparison: 3 vs 4Cα(p<0.0001), vs 4Cβ (p<0.0001), 4Cα vs 4Cβ (p<0.0001)). The next most abundant class of synapses, SSBm+, had the highest proportion in layer 4Cα (30%), with layer 3B having 26% and 4Cβ at 14% which was significantly lower than the proportion in 4Cα (Fig. 6A, Table 3, χ², p < 0.0001; Bonferroni comparison: 3 vs 4Cα (p=0.0083), vs 4Cβ (p<0.0001), 4Cα vs 4Cβ (p=0.0011)). Finally, MSBm+ were distributed such that layer 4Cβ had the highest density, followed by layer 3B and layer 4Cα with the lowest density (L4Cβ: 13%, L3B: 6%, L4Cα: 4; Fig. 6A, Table 3, χ², p < 0.0001; Bonferroni comparison: 3 vs 4Cα (p=0.0001), vs 4Cβ (p<0.0001), 4Cα vs 4Cβ (p=0.0001)). Boutons with multiple PSD’s but no mitochrondria (MSBm-) were rarely found in 4Cα or 4Cβ, and never were identified in layer 3B.

**Table 3:** Percentage of MSBm+ boutons identified in ESPina that established 2, 3, or more than 3 synapses per bouton. (χ², p = 0.003, for 2 synapses: post-hoc: 3B vs 4Cα (****), 3B vs 4Cβ (ns) and 4Cα vs 4Cβ (***); for 3 synapses: 3B vs 4Cα (ns), 3B and 4Cβ (****), 4Cα and 4Cβ **). **** p < 0.0001; *** p < 0.001; ** p < 0.01; * p < 0.05; ns p > 0.05

| Number synapses | Layer 3B<br>(N=34; 71 syn) | Layer 4Cα<br>(N=14; 34 syn) | Layer 4Cβ<br>(N=51; 135 syn) |
| --- | --- | --- | --- |
| 2 | 31 (91 %) | 8 (57 %) | 28 (55 %) |
| 3 | 3 (9 %) | 6 (43 %) | 17 (33 %) |
| >3 | 0 | 0 | 6 (12 %) |
| Average syn | 2.1 | 2.4 | 2.6 |

Overall, in all three layers, most boutons had a single PSD per bouton (the SSBm+ and SSBm-categories), they accounted for about 93.6, 95.7, and 86.9 % of all asymmetric synapses. (3, 4Cα, and 4Cβ, respectively, χ², p < 0.001; Bonferroni comparison: 3 vs 4Cα (p<0.0001), vs 4Cβ (p<0.0069), 4Cα vs 4Cβ (p<0.0001)). Considering boutons with mitochondria, when the single and multiple classes are combined the overall distribution across layers was similar in layer 3B and 4Cα (32.3%, 34.4%) and significantly larger than in 4Cβ (27.1%; χ², p =0.0007; Bonferroni comparison: 3 vs 4Cα (p<0.0001), vs 4Cβ (p=0.8512, n.s), 4Cα vs 4Cβ (p<0.0001)).

### Multisynaptic contacts

As the MSBm⁺ boutons establish two or more synapses, we assessed whether the number of contacts per MSBm⁺ bouton differed across layers. On average, MSBm⁺ boutons in layers 4Cβ and 4Cα established more synapses than MSBm⁺ boutons in layer 3B (4Cβ: 2.6 synapses/bouton, n = 51; 4Cα: 2.4 synapses/bouton, n = 14; 3B: 2.1 synapses/bouton, n = 34; Table 3). A global chi-square test indicated significant differences in the distribution of synapse numbers across layers (χ², p = 0.003). In layer 3B, the majority of MSBm⁺ boutons established two synapses (91%, n = 31), with the remaining 9% establishing three synapses (n = 3) (Fig. 6C). In layer 4Cα, there were fewer boutons establishing two synapses than in layer 3 (57%, n = 8), and the proportion of boutons establishing three synapses was significantly higher than in layer 3B (43%, n = 6). Neither in layer 3 nor 4Cα, we found boutons establishing more than 3 synapses. In layer 4Cβ, 55% (n = 28) of MSBm⁺ boutons established two synapses, 33% (n = 17) established three synapses, and 12% (n = 6) established more than three synapses (Fig. 6C, Table 3). Pairwise comparisons showed that the proportion of boutons establishing two synapses differed significantly between 3 vs 4Cα (p<0.0001) and 4Cα vs 4Cβ (p = 0.0001), but not between 3B vs 4Cβ (p = 0.6016, ns). For boutons establishing three synapses, significant differences were found between 3B and 4Cβ (p = 0.0001) and between 4Cα and 4Cβ (p = 0.0021), but not between 3B vs 4Cα (p = 0.2715, ns). Our results indicate that layer 4Cβ not only contains more MSBm⁺ boutons than 4Cα or 3B, but also that these boutons establish a greater number of synaptic contacts per bouton. Notably, we observed one MSBm⁺ bouton in layer 4Cβ forming as many as six synapses. Although MSBm⁺ boutons represent a minority of boutons in thalamocortical recipient layers, they can contribute a substantial fraction of the total synaptic input—accounting for up to approximately 15% of all synapses in layer 4Cβ.

### Postsynaptic targets

Next, we determined if each bouton category had a preferential postsynaptic target in the different layers. For each bouton we classified its postsynaptic element as dendrite (by the presence of mitochondria) or dendritic spine (by the presence of spine apparatus or lack of mitochondria); some cases were left unclassified when we were not able to observe the complete postsynaptic element.

When combining the three layers, asymmetric synapses preferentially target dendritic spines, 69 vs 31% for dendrites (Table 4, Fig 6B). The analysis of target distribution by layers showed that layer 3B had significantly more synapses on spines than the other layers (74%, 66%, and 66% for layer 3B, 4Cα, 4Cβ, respectively; Fig. 6B, Table 5, χ², p<0.0001, Bonferroni comparison: 3 vs 4Cα (p<0.0001), vs 4Cβ (p<0.0001), 4Cα vs 4Cβ (p<0.0001)).

**Table 4:**
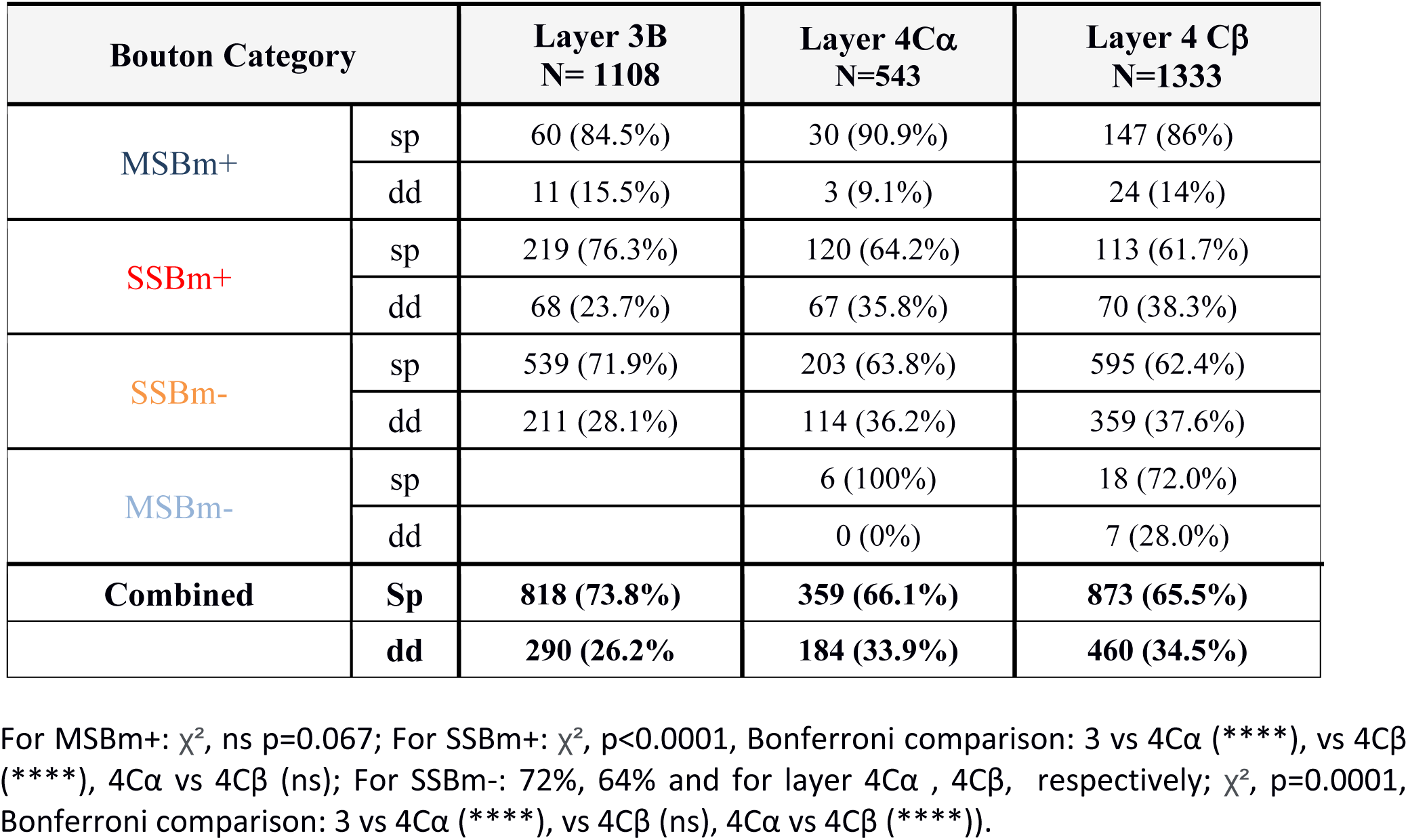
Percentage of the postsynaptic targets (dendrite or spine) associated with each bouton category in each layer. The last row (Combined) shows the data combined across all four categories. (χ², p<0.0001, Bonferroni comparison: 3 vs 4Cα (****), vs 4Cβ (****), 4Cα vs 4Cβ (****)). **** p < 0.0001; *** p < 0.001; ** p < 0.01; * p < 0.05; ns p > 0.05

**Table 5:** Percentage of the PSD morphology (macular, perforated, or horseshoe) associated with each synapse, for all layers combined (Total) and within each layer, only in the synapses that we were able to identify or were completed. (X^2^, p=0.089).

|  | **Total** | **3B** | **4Cα** | **4Cβ** |
| --- | --- | --- | --- | --- |
| Macular | 1661 (92%) | 656 (91.2%) | 370 (91.6%) | 635 (93.1%) |
| Perforated | 71 (4%) | 32 (4.5%) | 22 (5.4%) | 17 (2.5%) |
| Horseshoe | 73 (4%) | 31 (4.3%) | 12 (3%) | 30 (4.4%) |
| **total** | **1805** | **719** | **404** | **682** |

When we determined the type of postsynaptic target contacted by different bouton categories across layers we observed that MSBm+ boutons established significantly more contacts with dendritic spines than any other categories across all layers (Fig. 6B, Table 4, 86%, 69%, and 66%, for MSBm+, SSBm+, and SSBm-, respectively; χ², p<0.0001, Bonferroni comparison: 3 vs 4Cα (p<0.0001), vs 4Cβ (p<0.0001), 4Cα vs 4Cβ (p<0.0001)). However, we did not find differences in the percentage of contacts on spines made by MSBm+ by layer (χ², ns p=0.067) (85%, 91%, and 86%, for layer 3B, 4Cα, 4Cβ, respectively). However, both SSBm+ and SSBm-established more synapses in spines in layer 3B than in the other categories (Fig 6B, Table 4) (For SSBm+: 76%, 63%, and 62%, for layer 3B, 4Cα, 4Cβ, respectively; χ², p<0.0001, Bonferroni comparison: 3 vs 4Cα (p<0.0001), vs 4Cβ (p<0.0001), 4Cα vs 4Cβ (p=0.6131); For SSBm-: 72%, 64% and for layer 4Cα, 4Cβ, respectively; χ², p=0.0001, Bonferroni comparison: 3 vs 4Cα (p<0.0001), vs 4Cβ (p<0.499, ns), 4Cα vs 4Cβ (p<0.0001)).

#### PSD morphology

Synapses are characterized by a morphological and functional specialization of the postsynaptic membrane called the postsynaptic density (PSD). The PSDs exhibit diverse morphologies, that have been related to different numbers, proportion, and receptor expression (Nicholson & Geinisman, 2009). We classified the PSD in three commonly used shape categories: macular (with a flat, disk-shaped PSD), perforated (with one or more holes in the PSD), and horseshoe (with an indentation in the perimeter of the PSD) (Santuy et al. 2018; Domínguez-Álvaro et al. 2019). On average, macular are the most abundant in the three layers (92%), followed by perforated and horseshoe, each of the latter two representing 4% of the synapses (Figure 6D, Table 5). The distribution into shape category was not significantly different across layers (X^2^, p=0.089).

Next, we analyzed the different PSD morphologies associated with different boutons categories in the three layers.

In layer 3B, horseshoe and perforated were significantly more frequently associated with MSBm+ and SSBm+ bouton categories while macular morphology was significantly more frequently associated with SSBm-(Figure 6D, Table 6, χ², p<0.001, macular: Bonferroni comparison: MSBm+ vs SSBm+ (p<0.0001), vs SSBm- (p<0.0001), SSBm+ vs SSBm- (p<0.0001); horseshoe: Bonferroni comparison: MSBm+ vs SSBm+ (p=0.0111), vs SSBm- (p=0.0443), SSBm+ vs SSBm- (p=0.5600, ns)).; perforated: Bonferroni comparison: MSBm+ vs SSBm+ (p=0.281), vs SSBm- (p=0.0111), SSBm+ vs SSBm- (p<0.7027, ns)). In layer 4Cα, macular was the most abundant PSD morphology and was more frequently found in both SSBm+ and SSBm- than in MSBm+ (Figure 6D, Table 6, χ², p=0.0001; Bonferroni comparison: MSBm+ vs SSBm+ (p<0.0001), vs SSBm- (p<0.0001), SSBm+ vs SSBm- (p<0.0001)). Perforated synapses were more frequent associated with MSBm+ (Figure 6D, Table 6, χ², p=0.0001; Bonferroni comparison: MSBm+ vs SSBm+ (p=0.1290,ns), vs SSBm- (p=0.0310), SSBm+ vs SSBm- (p<0.4861, ns)). Horseshoe tends to be more frequently found in MSBm+ (Figure 6D, Table 6, χ², p<0.001, Bonferroni comparison: 3 vs 4Cα (p=0.4773, ns), vs 4Cβ (p=0.7375, ns), 4Cα vs 4Cβ (p=0.7042, ns)). In layer 4Cβ, macular was the most abundant PSD morphology for all the different categories (Figure 6D, Table 6, χ², p=0.0001; Bonferroni comparison: MSBm+ vs SSBm+ (p<0.0667), vs SSBm- (p<0.0001), SSBm+ vs SSBm- (p<0.0001)). The horseshoe morphology was most frequently associated with MSB+ boutons (Figure 6D, Table 6, χ², p<0.001, Bonferroni comparison: MSBm+ vs SSBm+(p=0.0016), vs SSBm- (p=0.0175), SSBm+ vs SSBm- (p=0.3638, ns)), while the perforated morphology was not significantly associated with any category (Figure 6D, Table 6, χ², p<0.001, Bonferroni comparison: MSBm+ vs SSBm+ (p=0.5256, ns), vs SSBm- (p=3638,ns), SSBm+ vs SSBm- (p=0.7805, ns))

**Table 6:**
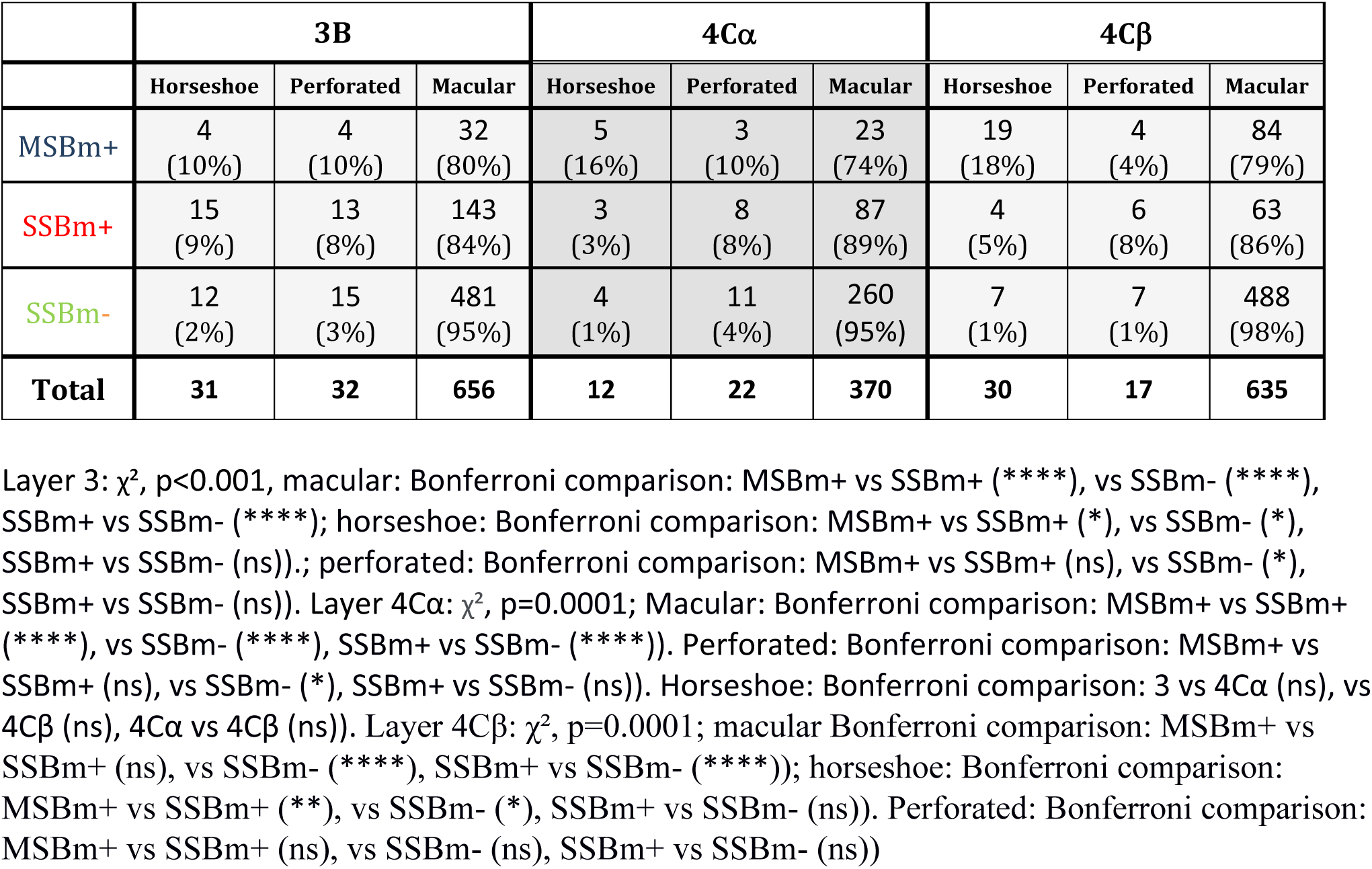
Percentage of the PSD morphology (macular, perforated, or horseshoe) associated with each bouton category and for each layer.

|  | **3B** |  |  | **4Cα** |  |  | **4Cβ** |  |  |
| --- | --- | --- | --- | --- | --- | --- | --- | --- | --- |
|  | Horseshoe | Perforated | Macular | Horseshoe | Perforated | Macular | Horseshoe | Perforated | Macular |
| MSBm+ | 4<br>(10%) | 4<br>(10%) | 32<br>(80%) | 5<br>(16%) | 3<br>(10%) | 23<br>(74%) | 19<br>(18%) | 4<br>(4%) | 84<br>(79%) |
| SSBm+ | 15<br>(9%) | 13<br>(8%) | 143<br>(84%) | 3<br>(3%) | 8<br>(8%) | 87<br>(89%) | 4<br>(5%) | 6<br>(8%) | 63<br>(86%) |
| SSBm- | 12<br>(2%) | 15<br>(3%) | 481<br>(95%) | 4<br>(1%) | 11<br>(4%) | 260<br>(95%) | 7<br>(1%) | 7<br>(1%) | 488<br>(98%) |
| **Total** | **31** | **32** | **656** | **12** | **22** | **370** | **30** | **17** | **635** |

Layer 3: $\chi^2$ , $p < 0.001$ , macular: Bonferroni comparison: MSBm+ vs SSBm+ (\*\*\*\*), vs SSBm- (\*\*\*\*), SSBm+ vs SSBm- (\*\*\*\*); horseshoe: Bonferroni comparison: MSBm+ vs SSBm+ (\*), vs SSBm- (\*), SSBm+ vs SSBm- (ns).; perforated: Bonferroni comparison: MSBm+ vs SSBm+ (ns), vs SSBm- (\*), SSBm+ vs SSBm- (ns)). Layer 4Cα : $\chi^2$ , $p = 0.0001$ ; Macular: Bonferroni comparison: MSBm+ vs SSBm+ (\*\*\*\*), vs SSBm- (\*\*\*\*), SSBm+ vs SSBm- (\*\*\*\*)). Perforated: Bonferroni comparison: MSBm+ vs SSBm+ (ns), vs SSBm- (\*), SSBm+ vs SSBm- (ns)). Horseshoe: Bonferroni comparison: 3 vs 4Cα (ns), vs 4Cβ (ns), 4Cα vs 4Cβ (ns)). Layer 4Cβ : $\chi^2$ , $p = 0.0001$ ; macular Bonferroni comparison: MSBm+ vs SSBm+ (ns), vs SSBm- (\*\*\*\*), SSBm+ vs SSBm- (\*\*\*\*)); horseshoe: Bonferroni comparison: MSBm+ vs SSBm+ (\*\*), vs SSBm- (\*), SSBm+ vs SSBm- (ns)). Perforated: Bonferroni comparison: MSBm+ vs SSBm+ (ns), vs SSBm- (ns), SSBm+ vs SSBm- (ns))

### 3D reconstructions of boutons in layers 3B, 4Cα, 4Cβ

To further extend the anatomical description of the different bouton categories we used the image stacks from the FIB/SEM to completely reconstruct boutons in layers 3B, 4Cα, and 4Cβ. We fully reconstructed 45, 48 and 63 boutons in layer 3B, 4Cα, 4Cβ, respectively. These full reconstructions enabled us to measure the volume of each bouton, the total volume of mitochondria inside the bouton, the number synaptic contacts per bouton, and importantly the area of the PSD. Using these measurements, we examined the relationships between bouton size, the number of synaptic contacts per bouton, the percentage of total volume occupied by mitochondria, and the area of the PSD, across the bouton categories in different layers.

#### Bouton size

The segmented synapses were assigned to different boutons categories based on the number of synapses that they established and the presence of mitochondria as described earlier. We found that on average, the bouton volumes across all categories, were significantly smaller in layer 3B than in 4Cα and 4Cβ (0.33, 0.96, and 1.09 µm^3^ for layers 3B (n = 45), 4Cα (n = 48), 4Cβ (n = 63), respectively, Fig. 7A; Table 8, KW, p<0.0001, 3 vs 4Cα (p=0.0012), 3 vs 4Cβ (p=0.0002), 4Cα vs 4Cβ (p=0.9567, ns). The MSBm+ synapses had significantly larger volumes (1.80 µm^3^) than SSBm+ (0.47 µm^3^) while SSBm- had the smallest volumes (0.15 µm^3^) (Fig. 7**B**; Table 8, KW, p<0.0001, MSBm+ vs SSBm+ (p<0.0001), MSBm+ vs SSBm- (p<0.0001), SSBm+ vs SSBm- (p<0.0001)). The MSBm+ boutons, which had the largest volumes, were similar in average volume in 4Cα (2.16 µm^3^) and 4Cβ (1.91 µm^3^) while the average MSB+ volume in layer 3B was considerably lower (0.66 µm^3^) (Table 8, Fig 7C). Although there was a clear difference in the volume among MSBm+ in three layers, the post-hoc comparison of the Kruskal-Wallis test did not show any significant difference in the median (KW<0.0001, post-hoc: MSBm+, 3 vs 4Cα (p=0.2810, ns), vs 4Cβ (p=0.1869, ns); 4Cα vs 4Cβ, (p=0.9999, ns), however, when the cumulative frequency distribution was tested, the Kolmogorov-Smirnov (K-S) testdetermined that the distribution of volumes of the MSB boutons in the TC recipient layers were significantly different from layer 3 (Fig. 7D-E; K-S: layer 3 vs 4Cα, p =0.0020, layer 3 vs 4Cβ, p =0.0055, layer 4Cα vs 4Cβ, p=0.7028, ns). For SSBm+ boutons, the largest were found in layer 4Cα (0.57 µm^3^), followed by 4Cβ (0.49 µm^3^), and the bouton volumes in both TC input layers were larger than SSBm+ in layer 3B (0.36 µm^3^), but there was no significant difference in these particular layers and categories (Table 8, Fig. 7C, KW< 0.0001, post-hoc: SSBm+, 3 vs 4Cα (p=0.5435, ns), vs 4Cβ (p=0.4501, ns); 4Cα vs 4Cβ, (p=0.9999, ns); again, the analysis of the cumulative frequency distribution showed that SSBm+ in 4Cα were significantly larger than in L3 (Fig. 7D-E; K-S: layer 3 vs 4Cα, p=0.0076; layer 3 vs 4Cβ,p=0.1008,ns; layer 4Cα vs 4Cβ, p=0.0916, ns). The smallest boutons are the SSBm- boutons that lack mitochondria and established only one synapse. There was no significance difference in either in the median bouton volume or the cumulative frequency distribution of these axon terminals in the different layers (0.13, 0.17, 0.17 µm^3^ for layers 3B, 4Cα, 4Cβ, respectively (Fig. 7D-E, Table 8, KW<0.0001; 3 vs 4Cα (p=0.3335, ns), vs 4Cβ (p=0.7291, ns); 4Cα vs 4Cβ, (p=0.9998, ns); K-S, layer 3 vs 4Cα, p=0.0414; layer 3 vs 4Cβ p=0.3208, ns; layer 4Cα vs 4Cβ,p=0.4889, ns).

**Figure 7:**
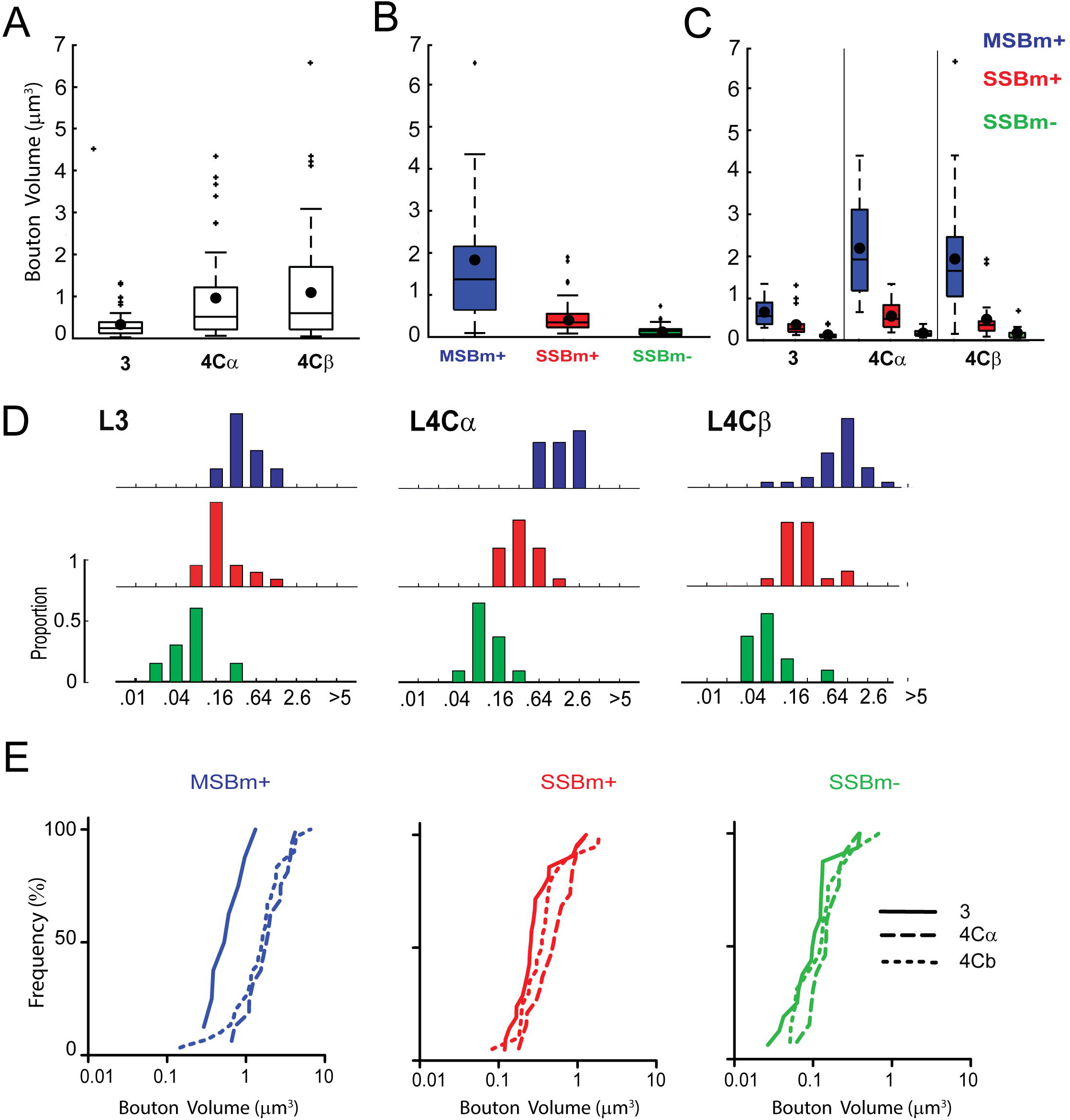
Distribution of bouton volume across categories and cortical layers. A) Boxplot showing the variation in the bouton volume in the three layers 3, 4Cα, and 4Cβ **(KW,** p<0.0001, post-hoc: 3 vs 4Cα (**), 3 vs 4CJ3 (***), 4Cα vs 4Cβ (ns). B) Boxplot showing the bouton volume variation by category (MSBm+ (blue), SSBm+ (red), and SSBm- (green) (KW, p<0.0001, post-hoc: MSBm+ vs SSBm+ (****), MSBm+ vs SSBm- (****, SSBm+ vs SSBm- (****). C) Boxplot showing the bouton volume variation by each layer (layers 3, 4Cα, and 4Cβ and categories (MSBm+, SSBm+, and SSBm) (KW, p<0.0001, posthoc: L3: SSBm- vs SSBm+ (ns), SSBm- vs MSBm+ (*), SSBm+ vs MSBm+ (ns); L4Cα: SSBm- vs SSBm+ (*), SSBm- vs MSBm+ (****), SSBm+ vs MSBm+ ns); L4CJ3: SSBm- vs SSBm+ (ns), SSBm- vs MSBm+ (****), SSBm+ vs MSBm+ (ns). D) Histograms showing the bouton volume distribution of each axon by layers 3, 4Cα, and 4Cβ and categories (MSBm+, SSBm+, and SSBm). E) Cumulative distributions of PSD’s areas by categories and layers for MSB (K-S: layer 3 vs 4Cα, p ***, layer 3 vs 4Cβ, ***, layer 4Cα vs 4Cβ, ns); for SSBm+ (K-S: layer 3 vs 4Cα, **; layer 3 vs 4Cβ, ns; layer 4Cα vs 4Cβ, ns), for SSBm- (K-S: layer 3 vs 4Cα, *; layer 3 vs 4Cβ, ns; layer 4Cα vs 4Cβ, ns). For A: Each box represents the interquartile range (IQR; 25th to 75th percentile), with the horizontal line inside the box indicating the median. The filled circle within each box denotes the mean bouton volume. Whiskers extend to the most extreme data points within 1.5 times the IQRfrom the box edges. Individual points beyond the whiskers are plotted asoutliers. **** p < 0.0001; *** p < 0.001; ** p < 0.01; * p < 0.05; ns p > 0.05.

### Mitochondria size

The results indicate that presence of mitochondria differentially increases bouton volume by layer. The 3D bouton reconstructions included the segmentation of mitochondria within each bouton. The volume occupied by mitochondria was almost 4 times larger in 4Cα and 4Cβ than layer 3B (Figure 8A, Table 8, KW, p=0.0001, post-hoc: 3 vs 4Cα, p=0.0002; 3 vs 4Cβ, p=0.0002, 4Cα vs 4Cβ, p=0.9644, ns). When mitochondria were present in boutons with multiple contacts (MSBm+) in the TC recipient layers the bouton volume was 10 fold greater than in non-mitochondrial containing synapses (12.7 and 11.2 times larger MSBm+ vs SSBm-, in 4Cα (2.16 vs 0.17 µm^3^) and 4Cβ (1.91 vs 0.17 µm^3^), respectively, (Fig 7C-D, Table 7, L4Cα: MSBm+ vs SSBm- (M-W, p<0.0001), L4Cβ: MSBm+ vs SSBm- (M-W, p<0.0001)). Whereas in the non-TC recipient layer 3B, this increase in the volume was 5 fold, when comparing MSBm+ vs SSBm- (0.66 vs 0.13 µm^3^) (Fig 7C-D, Table 8, M-W, p=0.011). For boutons making only a single contact that had mitochondria the volume was 2.9 times greater than in non-mitochondrial containing synapses SSBm+ vs SSBm- in layer 3B (0.36 vs 0.13 µm^3^) and in 4Cβ (0.49 vs 0.17 µm^3^) (Fig 7C-D, Table 8, L3B: M-W, p=0.1792, ns; L4Cβ: M-W, p=0.1804, ns), and up to 3.4 times greater in 4cα (0.57 vs 0.17 µm^3^) (Table 8, L4Cα: M-W, p = 0.0499).

**Figure 8:**
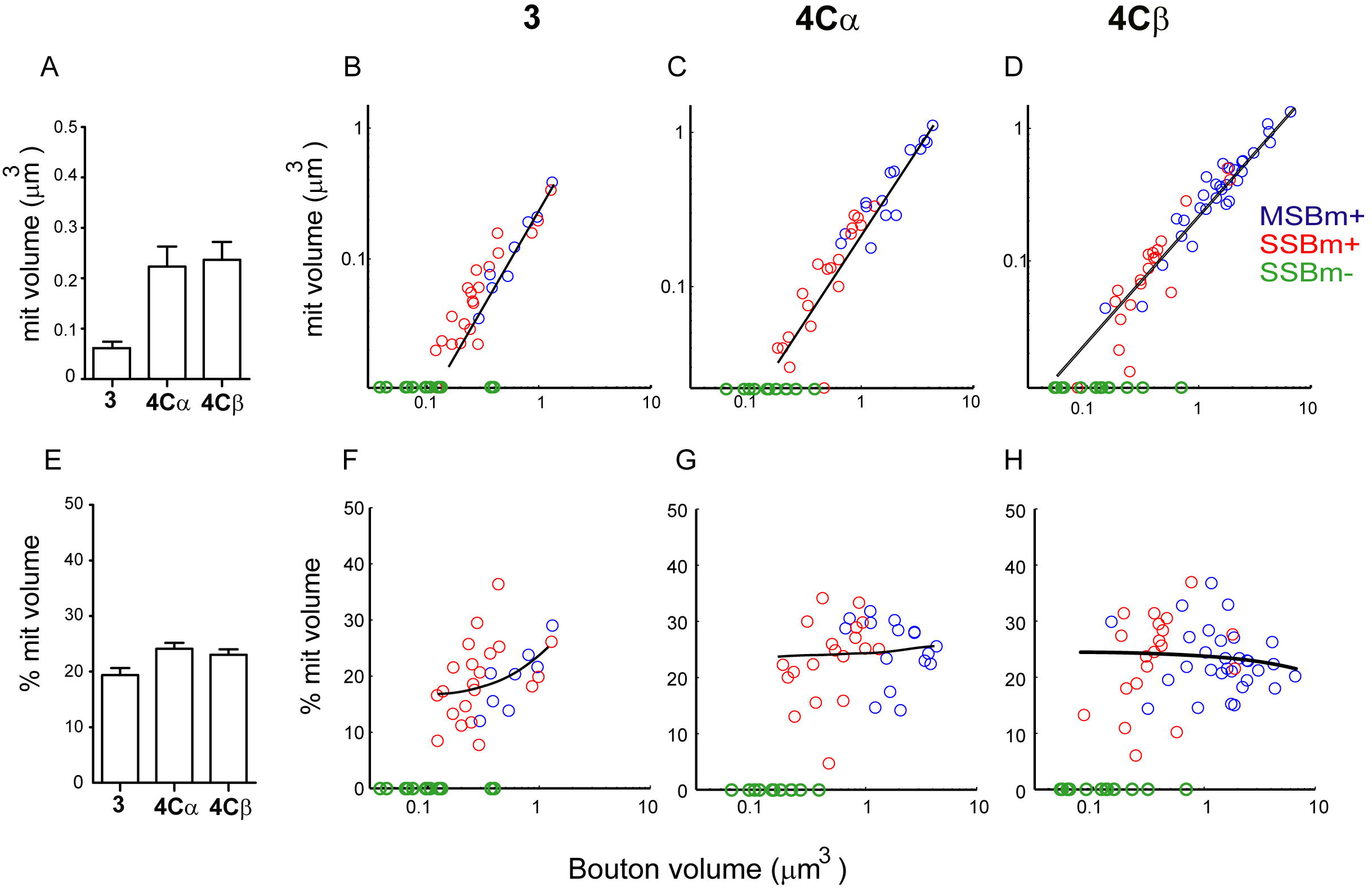
Relationship between bo11ton volume and mitochondrial volume inside each terminal. A) Graph showing the mean mitochondria volume inside each terminal in the three layers 3, 4Cα, and 4Cβ (KW, p=0.0001, post-hoc: 3 vs 4Cα, ***; 3 vs 4Cβ, ***, 4Cα vs 4Cβ, ns). B-D) Scatter plots showing a positive correlation between mitochondria volume and bouton volume for each bouton category in each layer [layer 3 (B, Spearman r==0.8540, ****), layer 4Cα (C, Spearman r==0.9404, ****), and layer 4CP (D, Spearman r==0.9259, ****)]. E) Graph showing the percentage of the axonal volu1ne occupied by mitochondria in the three layers 3, 4Cα, and 4Cβ KW, p=0.0065, post-hoc: 3 vs 4Cα, **; 3 vs 4Cβ, *; 4Cα vs 4Cβ, ns). F-H) Scatter plots showing the correlation between the percentage of 1nitochondria and bouton volume. In layers 4Cα (G, Spearman *r* = 0.1660, n.s.) and 4Cβ (H, Spearman *r* = -0.0855, n.s.), the percentage of mitochondria remains relatively constant across a tenfold range of bouton volumes. In contrast, layer 3 (F) shows a modest but significant positive correlation (Spearman *r* = 0.4712, **). **** p < 0.0001; *** p < 0.001; ** p < 0.01; * p < 0.05; ns p > 0.05

**Table 7:** Mean (± 1 SEM) axonal bouton volume (µm^3^) in each category and layer, and across layers (column Average) (KW, p<0.0001, 3 vs 4Cα (**), 3 vs 4Cβ (***), 4Cα vs 4Cβ (ns). and categories (row Average) KW, p<0.0001, MSBm+ vs SSBm+ (****), MSBm+ vs SSBm- (****), SSBm+ vs SSBm- (****).

| | 3B | 4Cα | 4Cβ | Average |
| --- | --- | --- | --- | --- |
| **MSBm+** | Mean $0.66 \pm 0.13$<br>Median 0.57<br>(N=8)<br>7 less than $1 \mu\text{m}^3$ (87%) | Mean $2.16 \pm 0.29$<br>Median 1.90<br>(N=16)<br>2 less than $1 \mu\text{m}^3$ (12%) | Mean $1.91 \pm 0.26$<br>Median 1.63<br>(N=30)<br>7 less than $1 \mu\text{m}^3$ (23%) | Mean $1.80 \pm 0.18$<br>Median 1.55<br>(N=54)<br>16 less than $1 \mu\text{m}^3$ (30%) |
| **SSBm+** | Mean $0.36 \pm 0.07$<br>Median 0.26<br>(N=21)<br>20 less than $1 \mu\text{m}^3$ (95%) | Mean $0.57 \pm 0.07$<br>Median 0.50<br>(N=19)<br>18 less than $1 \mu\text{m}^3$ (95%) | Mean $0.49 \pm 0.11$<br>Median 0.36<br>(N=20)<br>18 less than $1 \mu\text{m}^3$ (90%) | Mean $0.47 \pm 0.04$<br>Median 0.35<br>(N=60)<br>56 less than $1 \mu\text{m}^3$ (93%) |
| **SSBm-** | Mean $0.13 \pm 0.03$<br>Median 0.10<br>(N=16)<br>16 less than $1 \mu\text{m}^3$ (100%) | Mean $0.17 \pm 0.02$<br>Median 0.15<br>(N=13)<br>13 less than $1 \mu\text{m}^3$ (100%) | Mean $0.17 \pm 0.05$<br>Median 0.13<br>(N=13)<br>13 less than $1 \mu\text{m}^3$ (100%) | Mean $0.15 \pm 0.02$<br>Median 0.12<br>(N=42)<br>42 less than $1 \mu\text{m}^3$ (100%) |
| **Average** | Mean $0.33 \pm 0.05$<br>Median 0.25<br>(N=45)<br>43 less than $1 \mu\text{m}^3$ (96%) | Mean $0.96 \pm 0.15$<br>Median 0.52<br>(N=48)<br>33 less than $1 \mu\text{m}^3$ (69%) | Mean $1.09 \pm 0.16$<br>Median 0.60<br>(N=63)<br>38 less than $1 \mu\text{m}^3$ (60%) | |

**Table 8:** Mean (± 1 SEM) mitochondria volume (µm^3^) within each bouton in each layer KW, p=0.0001, post-hoc: 3 vs 4Cα, ***; 3 vs 4Cβ, ***, 4Cα vs 4Cβ, ns). and percentage (%) of bouton volume occupied by mitochondria in each layer Anova, p=0.0145, post-hoc: 3 vs 4Cα, *; 3 vs 4Cβ, *; 4Cα vs 4Cβ, ns).

|  | **3B** | **4Cα** | **4Cβ** |
| --- | --- | --- | --- |
| Mitochondria Volume<br>$\mu\text{m}^3$ | $0.0952 \pm 0.0116$<br>(N=29) | $0.3189 \pm 0.0359$<br>(N=35) | $0.3032 \pm 0.0357$<br>(N=50) |
| Mitochondria % | $19.41 \pm 0.83$<br>(N=29) | $24.09 \pm 0.82$<br>(N=35) | $23.07 \pm 0.83$<br>(N=50) |

The 3D morphology of the mitochondria in the reconstructed terminals was elongated (Fig 3B-C & Fig 12), some of them in a coil, concentrated in the terminal and occupying on average 19%, 24% and 23% of the bouton volume in layers 3B, 4Cα and 4Cβ, respectively (Fig. 8E; Table 9, Anova, p=0.0145; post-hoc: 3 vs 4Cα, p=0.0151; 3 vs 4Cβ, p=0.0493; 4Cα vs 4Cβ, p=0.7624, ns).

**Table 9:** Mean (± 1 SEM) PSD area (µm^2^) associated with individual synapses for each bouton category and layers. Averages as in Table 7. (KW, p<0.0001, post-hoc: 3 vs 4Cα, **, 3 vs 4Cβ, ****; 4Cα vs 4Cβ, ns)

|  | **3B** | **4Cα** | **4Cβ** | **Average** |
| --- | --- | --- | --- | --- |
| **MSBm+** | Mean $0.109 \pm 0.01$<br>Median 0.095<br>(N=17) | Mean $0.165 \pm 0.02$<br>Median 0.154<br>(N=36) | Mean $0.175 \pm 0.01$<br>Median 0.144<br>(N=87) | Mean $0.164 \pm 0.01$<br>Median 0.138<br>(N=140) |
| **SSBm+** | Mean $0.115 \pm 0.01$<br>Median 0.093<br>(N=21) | Mean $0.171 \pm 0.02$<br>Median 0.147<br>(N=19) | Mean $0.160 \pm 0.01$<br>Median 0.151<br>(N=20) | Mean $0.148 \pm 0.01$<br>Median 0.140<br>(N=60) |
| **SSBm-** | Mean $0.090 \pm 0.01$<br>Median 0.072<br>(N=16) | Mean $0.148 \pm 0.02$<br>Median 0.107<br>(N=13) | Mean $0.147 \pm 0.03$<br>Median 0.128<br>(N=13) | Mean $0.126 \pm 0.08$<br>Median 0.103<br>(N=42) |
| **Average** | Mean $0.106 \pm 0.01$<br>Median 0.088<br>(N=54) | Mean $0.157 \pm 0.01$<br>Median 0.143<br>(N=72) | Mean $0.169 \pm 0.01$<br>Median 0.142<br>(N=123) | |

Our results we demonstrate that bouton volume scales with the mitochondrial volume (Fig 8B-D), as we found a strong positive correlation between the bouton volume and the mitochondria volume (Fig. 8B-D) in the two axonal categories that have mitochondria (MSBm+ and SSBm+) and in all the layers (layer 3B: Spearman r=0.8540, p<0.0001; layer 4Cα: Spearman r=0.9404, p<0.0001; layer 4Cβ:Spearman r=0.9259, p<0.0001). However, the proportion of the bouton occupied by the mitochondria remains constant at 22% of the total volume in layer 4Cα and 4Cβ (Fig 8F-H, Table 8; layer 4Cα: Spearman r=0.1660, p=0.5950 ns; layer 4Cβ: Spearman r=-0.0855, p=0.6102 ns). Only in layer 3B was there a positive relationship between the increase of the volume and the percentage of the volume occupied by mitochondria (Fig 8F, Table 8; Spearman r=0.4712, p<0.0099).

The findings above raised the question of whether the difference in volume between boutons containing mitochondria and those without was solely due to the presence of mitochondria or if there was an additional non-mitochondrial volume increase associated with the MSBm+ and SSBm+ boutons. To address this, we compared the volume of SSBm-terminals with the volume of SSBm+ terminals, subtracting the mitochondrial volume. Our analysis revealed that the bouton volume, excluding mitochondria, of the SSBm+ terminals was significantly larger than that of the SSBm-terminals across all three layers [Layer 3B: 0.1257 vs. 0.2853 µm³, SSBm- vs. SSBm+ without mitochondria (M-W, p = 0.0004); Layer 4Cα: 0.1652 vs. 0.4284 µm³, SSBm- vs. SSBm+ without mitochondria (M-W, p = 0.0002); Layer 4Cβ: 0.1716 vs. 0.3685 µm³, SSBm- vs. SSBm+ without mitochondria (M-W, p = 0.0043) (Supplementary Figure 3)]. These results suggest that the increase in volume of the SSBm+ terminals cannot be solely attributed to the volume of the mitochondria, suggesting more local cytoplasm that could potentially accommodate more synaptic vesicles and the machinery for neurotransmitter release.

### PSD number and size

To test the hypothesis that boutons containing mitochondria could potentially accommodate more synaptic vesicles and neurotransmitter release machinery, and consequently establish more synapses, we investigated the relationship between the number of synapses each bouton formed and the bouton volume across categories and layers (Fig. 9). For all the 3D-reconstructed axons across the three layers, we found a positive correlation between bouton volume and the number of synapses (Fig. 9A). As bouton volume increased, the number of synapses also increased. Comparing the relationship between bouton volume and the number of PSDs across layers, we observed a positive correlation in all cases. However, this correlation was marginally stronger in the thalamocortical (TC) recipient layers: Spearman r = 0.5271, p = 0.0003 for Layer 3B; Spearman r = 0.7192, p < 0.0001 for Layer 4Cα; and Spearman r = 0.7825, p < 0.0001 for Layer 4Cβ (Fig. 9A–C). Interestingly, in layer 3B the vast majority (96%) of the boutons sampled were smaller than 1 µm^3^ (n=43, Fig. 9B), irrespective of the number of PSDs. In the TC recipient layers, the majority of boutons with a single PSD were smaller than 1 µm³; 100% of SSBm-boutons in both layers and 95% and 90% of SSBm+ boutons in 4Cα and 4Cβ, respectively. In contrast, 88% of the MSBm+ boutons in 4Cα and 77% in 4Cβ were larger than 1 µm³. These results show that the increase in the number of PSDs is associated with an increase in the bouton volume with an enhanced association in the TC recipient layers

**Figure 9:**
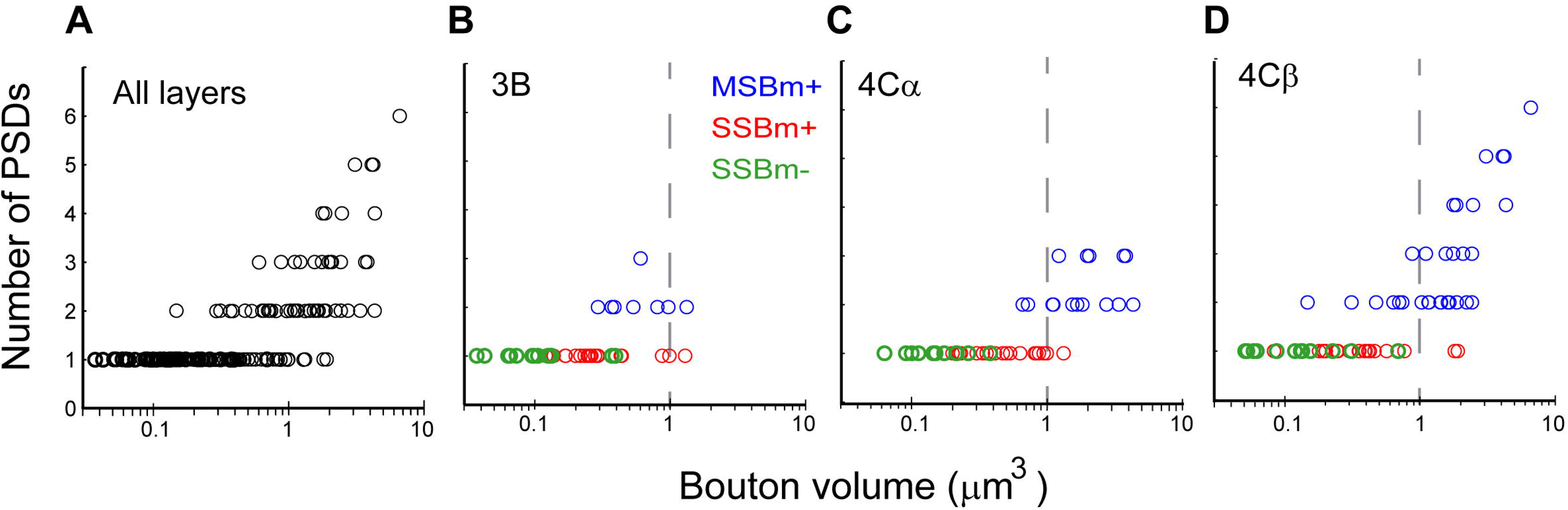
Distribution of number of PSD’s with bouton volume across layers and bouton category. A) Scatter plots showing a positive correlation between bouton volume and the number of PSDs across all layers. B) In layer 3B, a modest but statistically significant positive correlation is observed between bouton volume and the number of PSDs (Spearman r == 0.5171, ***). C) In layer 4Cα, a strong positive correlation exists between bouton volume and the number of PSDs (Spearman r == 0.7192, ****). D) Similarly, a strong correlation is seen in layer 4Cβ (Spearman r == 0.7825, ****). The d.ashed line indicates a volume cutoff of 1 µm^3^, and the colors represent different bouton categories. **** p < 0.0001; *** p < 0.001; ** p < 0.01; * p < 0.05; ns p > 0.05

As the number of PSDs significantly increased with volume, we asked whether the area of individual PSDs also varied by layer or axonal category. We found that on average the PSD’s areas were significantly larger in 4Cα and 4Cβ than in layer 3B (0.106, 0.157, and 0.169 µm^2^ for layers 3B, 4Cα, 4Cβ, respectively, Fig. 10A; Table 9, (KW, p<0.0001, post-hoc: 3 vs 4Cα, p=0.0013, 3 vs 4Cβ, p<0.0001; 4Cα vs 4Cβ, p=0.8755, ns). When we compared the PSD area across axonal categories, we found that MSBm+ boutons had largest PSD’s with the largest area (0.1643 µm^2^), followed by SSBm+ (0.1476 µm^2^), and the smallest PSD’s areas were found in SSBm- (0.1255 µm^2^) however none of these differences were significant (Fig. 10B, Table 9, KW, p=0.0606, ns). Next, when the comparison of PSD’s areas by layer and category was made it was found that the largest PSD’s areas were in the MSBm+ boutons of layer 4C (0.1749 µm^2^ and 0.1649 µm^2^, for 4Cα and 4Cβ, respectively) and SSBm+ boutons in layer 4Cα (0.1706 µm^2^) and 4Cβ (0.1599 µm^2^) but none of these differences were significant (Fig. 10C-D, KW, p, ns). Although the PSD areas of boutons belonging to the MSBm+ and SSBm+ categories in the TC recipient layers were larger than their counterparts in layer 3B MSBm+ (0.1088 µm^2^) and SSBm+ (0.1152 µm^2^) the differences in area were not significant (Fig. 10C-D, Table 9). A cumulative frequency distribution analysis showed that the PSD area of MSBm+ in layer 3 was significantly different from the MSBm+ in 4Cα and 4Cβ (K-S: L3 vs 4Cα, p=0.0043, L3 vs 4Cβ, p=0.0023; 4Cα vs 4Cβ, p=0.7018, ns). (Fig, 10E). The boutons without mitochondria SSBm- had the smallest PSD’s areas in all the layers (0.090, 0.148, and 0.147 µm^2^ for layers 3B, 4Cα, 4Cβ, respectively (Fig. 10C-D, Table 9). A cumulative distribution analysis showed that PSD area of SSBm- was smaller in layer 3 than in the other layers ((K-S: L3 vs 4Cα, p=0.016, L3 vs 4Cβ, p=0.016; 4Cα vs 4Cβ, p=0.8281, n.s). A cumulative frequency distribution analysis showed that the PSD area of SSBm+ showed no significant differences among the three categories (K-S: L3 vs 4Cα, p=0.0513 ns, L3 vs 4Cβ, p=0.0794, ns; 4Cα vs 4Cβ, p=0.8355, ns). (Fig, 10E). These results show that PSD area are larger in TC recipient layers than L3 particularly in MSB+ and SSBm- categories.

**Figure 10:**
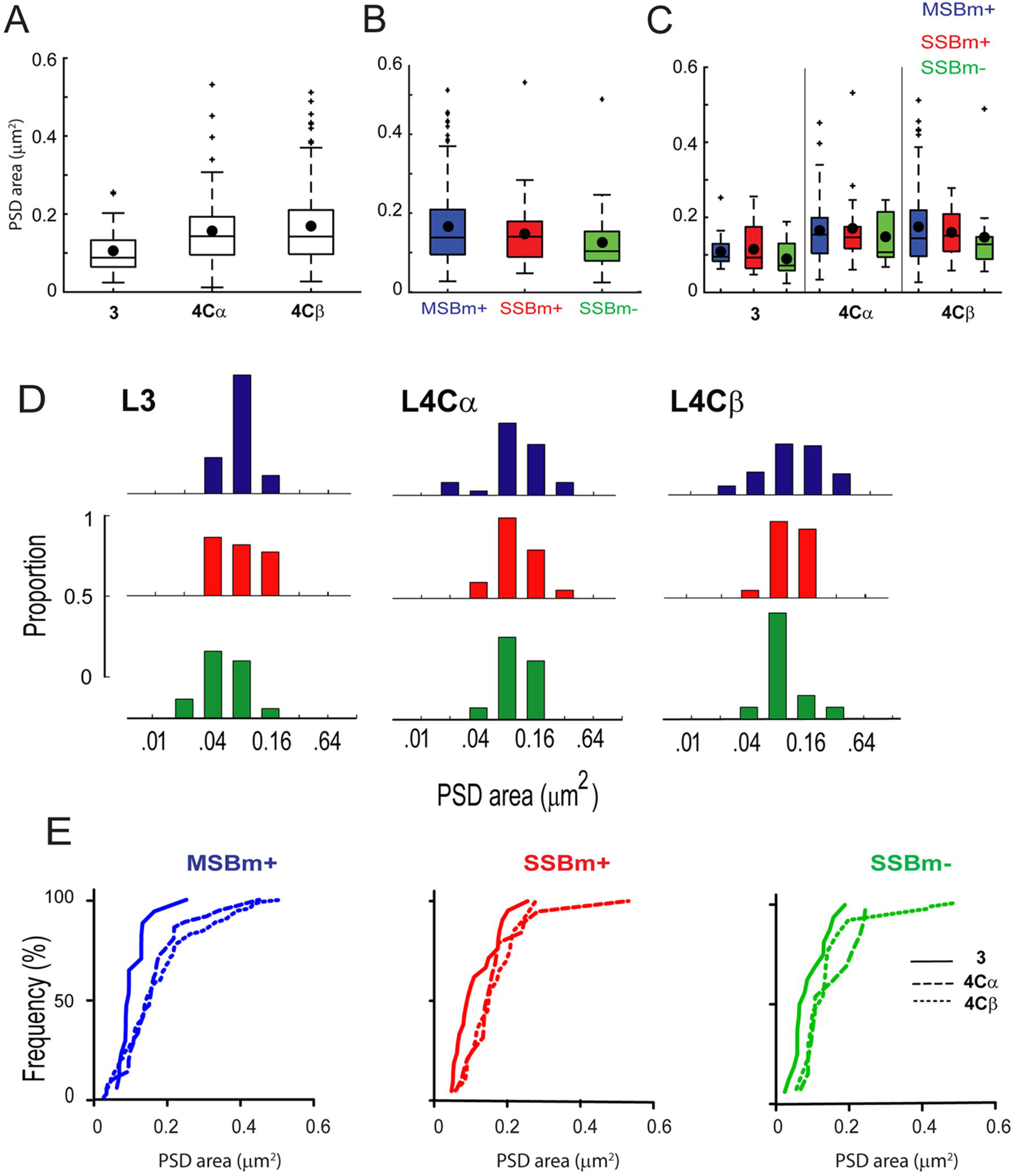
Distribution of PSD area for different bouton categories and across cortical layers. A) Boxplots showing the variation in the PSD in the three layers 3, 4Cα, and 4Cβ(KW, p<0.0001, post-hoc: 3 vs 4Cα, **, 3 vs 4Cβ, ****; 4Cα vs 4Cβ, ns). B) Boxplots showing the PSD area variation for each bouton category (MSBm+ (blue), SSBm+ (red), and SSBm- (green) (KW, p=0.0616, ns). C) Boxplots showing PSD area distribution for each layer (layers 3, 4Cα, and 4Cβ and category (MSBm+, SSBm+, and SSBm) (KW, 0.018; post-hoc: L3: SSBm- vs SSBm+ (ns), SSBm- vs MSBm+ (p=0.9973, ns), SSBm+ vs MSBm+ (ns); L4Cα: SSBm- vs SSBm+ (ns), SSBm- vs MSBm+ (, ns), SSBm+ vs MSBm+ (ns); L4Cb: SSBm- vs SSBm+ (ns), SSBm- vs MSBm+ (ns), SSBm+ vs MSBm+ (p=0.9999, ns). D) Histograms showing the PSD area distribution of boutons by layer (3, 4Cα, and 4Cβ and category (MSBm+, SSBm+, and SSBm). E) Cumulative distributions of PSD’s areas by categories and layers for MSB (K-S: L3 vs 4Cα, **, L3 vs 4Cβ, **; 4Cα vs 4Cβ, ns); for SSBm+ (K-S: L3 vs 4Cα, ns, L3 vs 4Cβ, ns; 4Cα vs 4Cβ, ns); for SSBm- (K-S: L3 vs 4Cα, *, L3 vs 4Cβ, *; 4Cα vs 4Cβ, n.s). For A: Each box represents the interquartile range (IQR; 25th to 75th percentile), with the horizontal line inside the box indicating the median. The filled circle within each box denotes the mean bouton volume. Whiskers extend to the most extreme data points within 1.5 times the IQR from the box edges. Individual points beyond the whiskers are plotted as outliers. **** p < 0.0001; *** p < 0.001; ** p < 0.01; * p < 0.05; ns p > 0.05 MSBm+ (p=0.9973, ns), SSBm+ vs MSBm+ (ns); L4Cα: SSBm-vs SSBm+ (ns), SSBm-vs MSBm+ (, ns), SSBm+ vs MSBm+ (ns); L4Cb: SSBm- vs SSBm+ (ns), SSBm- vs MSBm+ (ns), SSBm+ vs MSBm+ (p=0.9999, ns). D) Histograms showing the PSD area distribution of boutons by layer (3, 4Cα, and 4Cβ and category (MSBm+, SSBm+, and SSBm). E) Cumulative distributions of PSD’s areas by categories and layers for MSB (K-S: L3 vs 4Cα, **, L3 vs 4Cβ, **; 4Cα vs 4Cβ, ns); for SSBm+ (K-S: L3 vs 4Cα, ns, L3 vs 4Cβ, ns; 4Cα vs 4Cβ, ns); for SSBm- (K-S: L3 vs 4Cα, *, L3 vs 4Cβ, *; 4Cα vs 4Cβ, n.s). For A: Each box represents the interquartile range (IQR; 25th to 75th percentile), with the horizontal line inside the box indicating the median. The filled circle within each box denotes the mean bouton volume. Whiskers extend to the most extreme data points within 1.5 times the IQR from the box edges. Individual points beyond the whiskers are plotted as outliers. **** p < 0.0001; *** p < 0.001; ** p < 0.01; * p < 0.05; ns p > 0.05

To complement these results, we determined the relationship between the total PSD area and bouton volume after subtracting mitochondrial volume. This adjustment allowed us to compare bouton volumes independently of mitochondrial contribution. For all the boutons in all layers there was a strong correlation between the summed PSD area and the bouton volume independent of the mitochondria (Fig. 11A, Slope: 0.5032 ; R-squared: 0.5286; p-value: <0.0001). The same relationship betweenaxonal volume and PSD area was observed in individual bouton categories. For MSBm+ (Fig. 11B; Slope: 0.5407, Intercept: -1.1006, R-squared: 0.4024, p-value:<0.0001),] SSBm+ (Fig. 11C; Slope: 0.3602, Intercept: -1.5889, R-squared: 0.2110, p-value: 0.0002 and SSBm-, (Fig. 11D; Slope: 0.5075, Intercept: -1.1591, R-squared: 0.3656, p-value: <0.0001) there was a similar strong correlation between bouton volume (minus mitochrondrial volume) and summed PSD area. These results suggest that while there is a consistent trend across synaptic boutons, the presence of multiple synapses might be associated with a stronger and more consistent correlation between axonal volume and PSD measures

**Figure 11:**
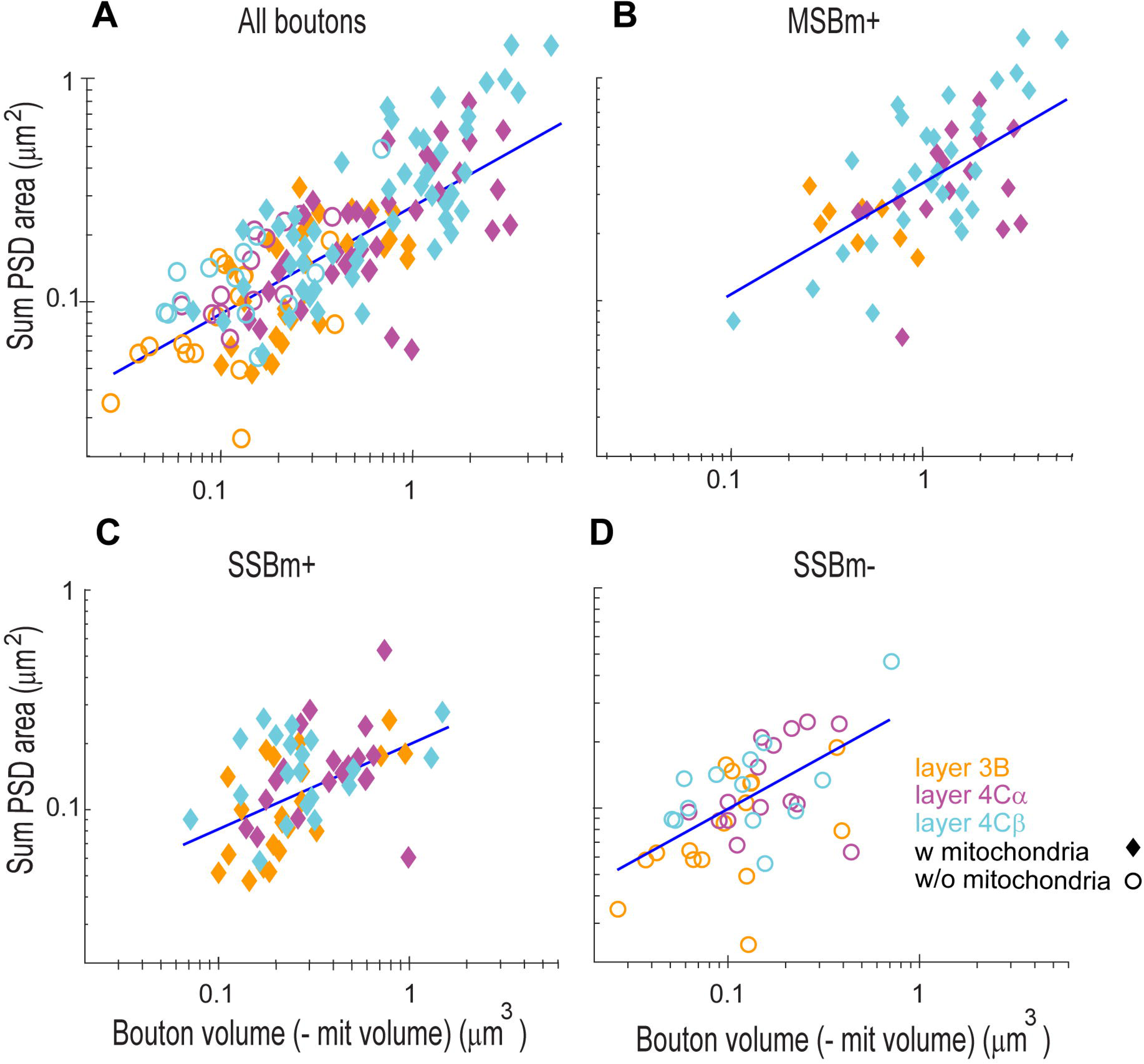
Relationship between bouton volume and PSD area for different boutons categories and layers. A) Scatter plots showing a positive correlation between the bouto11 volu1ne minus mitochondrial volume and the sum of all PSDs made by each bouton in tl1e three layers. (Slope: 0.5032; Intercept: - 1.2368; R-squared: 0.5286; ****). B) For MSB+, the bouton volume minus the mitochondria volume also shows a positive correlation with the PSD area (Slope: 0.5407, Intercept: -1.1006, R-squared: 0.4024, ****). C) For SSBm+, the bouton volume minus the mitochondria volu1ne also shows a positive correlation with the PSD area (Slope: 0.3602, Intercept: -1.5889, R-squared: 0.2110, ***).D) For boutons without mitochondria, SSBm-, the bouton volume also shows a positive correlation with the PSD area (Slope: 0.5075, Intercept: -1.1591, R-squared: 0.3656, ****). ♦, indicates boutons with mitochondria, o, indicates boutons without mitochondria.**** p < 0.0001; *** p < 0.001; ** p < 0.01; * p < 0.05; ns p > 0.05

### Other axonal morphologies

In addition to the bouton categories described above, we observed two examples of thin *en passant* fibers in layer 4Cβ that did not contain an obvious varicosal organization (Figure 12 I-J, compared to the MSBm+ Fig12 A-H), yet were rich in mitochondria and had multiple clustered PSDs. The first example (Fig. 12I axon 18), had a length of 13.06 µm, a mean diameter of 0.88 ± 0.32 µm (mean ± 1SD; range 0.1-1.3 µm), established 12 asymmetric synapses, 9 on spines and 3 on dendrites. This axon had one cluster of 7 synapses in a small region of 1.9 x 0.76 x 0.99 = 1.6 µm^3^ (1.67 µm^3^ after shrinkage) and the mitochondria were not throughout the length of the axon segment but were located in the region where the PSDs were clustered and close to the other synapses. The second example (Fig 11J, axon 8) had a length of 12.2 µm, a diameter of 0.91 ± 0.25 µm (mean ± 1SD; range 0.4-1.4 µm), established 7 asymmetric synapses, all of them on spines, had one cluster of 3 synapses in a small region of 1.5 x 0.88 x 1.1 = 1.4 µm^3^ (1.6 µm^3^ after shrinkage) and mitochondria extended through the whole length of the axon segment. These observations suggest that en passant axons in layer 4Cβ, despite lacking obvious varicosal organization, exhibit a highly specialized architecture with rich mitochondrial content and spatially clustered synapses, highlighting their potential role in supporting localized synaptic activity and energy demands within this cortical layer.

**Figure 12:**
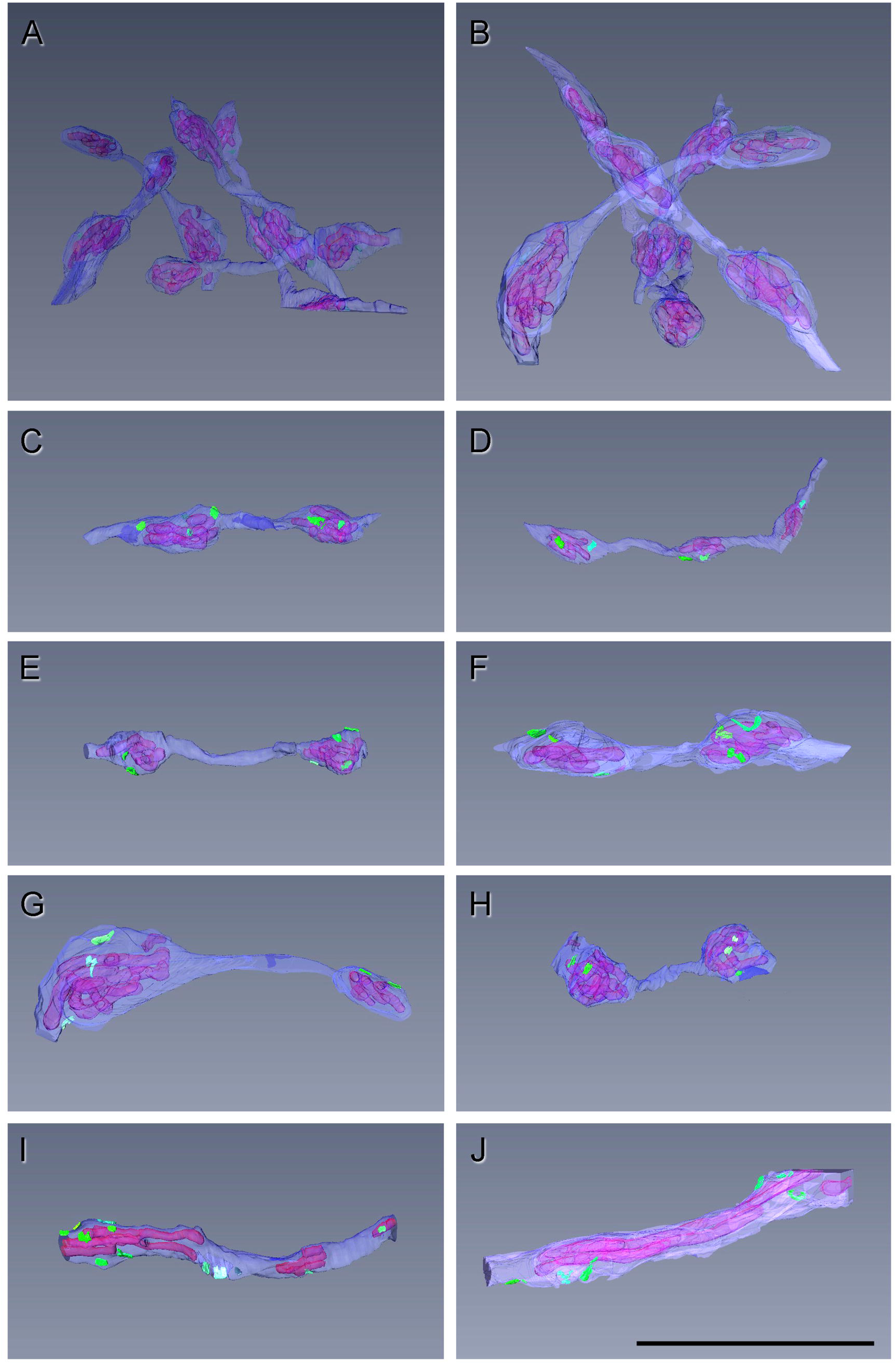
Three dimensional reconstructions of MSBm+ boutons showing their structural organization. A-B) Examples of MSBm+ boutons (blue) drawn in two different stacks of layer 4Cβ of macaque Vl. C-H) Examples of MSBm + with en passant morphology, boutons establishing synapses (greens) are filled with mitochondria (purple) and are linked together by thin axonal strand. 1-J) Examples of other axons establishing multiple synapses, with mitochondria but without the enlargement of the boutons. Synapses in green, mitochondria in transparent pink inside the boutons.

## DISCUSSION

The current study reveals important characteristics in the synaptic organization of different layers in macaque V1. Thalamic recipient layers (4Cα and 4Cβ) share with non-preferential TC recipient layers (layer 3B) a similar Excitatory/Inhibitory synaptic ratio, a similar proportion of postsynaptic targets, and a similar axonal morphology. However, the three layers also show important differences in the synaptic density, in the percentage of the different categories of axonal boutons, in bouton volume, PSD area, and total PSD area per bouton. These results suggest that each layer has a unique ultrastructural composition of synapses related to their position in the processing circuit.

### Synaptic density and E/I balance

Determining the density and morphology of axons in different brain layers is crucial to understand brain function. Earlier studies using 2D electron microscopy (2D-EM) reported synaptic densities in the same layers examined here. In layer 3B, O’Kusky & Colonnier (1982) found 0.34 synapses/µm³, while in layer 4C, densities ranged from 0.21–0.25 synapses/µm³ in 4Cα and 0.30–0.31 synapses/µm³ in 4Cβ (O’Kusky & Colonnier, 1982; Latawiec et al, 2000). In contrast, Beaulieu et al. (1992), also using 2D-EM, reported higher densities of 0.54 synapses/µm³ in layer 3B and 0.45 synapses/µm³ in layer 4C. However, the estimation of synaptic density particularly in 2D-EM is problematic when the profiles are not en face (DeFelipe et al. 1999; Kubota et al. 2009). More recent studies, including our own, suggest that some earlier 2D-EM studies likely underestimated synaptic density in layers 4Cα and 4Cβ of macaque V1 by nearly 50% (Medalla & Luebcke, 2015; Hsu et al. 2017; Garcia-Marin et al. 2019). Our new 3D-EM-based estimates of the synaptic density in V1 macaque layer 3B align well with those reported in rhesus V1 (0.58-0.64 synapses μm^3^; Medalla & Luebcke, 2015; Hsu et al. 2017) and with our previous findings in 4Cα and 4Cβ (Garcia-Marin et al. 2019).These findings highlight the importance of volumetric imaging for accurate quantification of synaptic architecture.

In contrast to the synaptic density changes between layers, our data showed that there is a constant ratio of asymmetric/symmetric synapses in the three layers with an average of 86 % vs 14% respectively. Previous results showed a similar constant ratio of asymmetric/symmetric synapses in layer 3B of primate V1, (84-86% asymmetric synapses vs 14-17% symmetric synapses) (Beaulieu et al.1992; Medalla & Luebcke, 2015; Hsu et al. 2017), and in layers 4Cα (89% vs 11%) and 4Cβ (88% vs 12%) (Latawiec et al. 2000). The asymmetric/symmetric synapse ratios in layers 4Cα and 4Cβ closely match the ratios of excitatory to inhibitory neurons in these layers (Kelly et al. 2019). This consistent E/I ratio suggests that inhibitory balance is a tightly regulated feature of local V1 circuitry, possibly critical for maintaining stable network function across cortical layers.

### Synaptic bouton categories and layer specialization

We examined whether the ultrastructural characteristics of the neuropil varied across layers and found distinct layer-specific distributions of synaptic bouton categories. Axonal bouton profiles were classified into four groups based on the number of synapses and the presence of mitochondria. The MSBm− category appeared exclusively in layer 4Cβ and represented a very small proportion of synapses. For the remainder of the analysis, we focused on the SSBm−, SSBm+, and MSBm+ categories.

Across all layers, the largest proportion consisted of SSBm− boutons, which were consistent across layers at approximately 72%. In contrast, the proportion of MSBm+ boutons proportion varied across layers. In layer 3B, our reported MSBm+ proportion was comparable to that observed in rhesus V1 by Hsu et al. (2017). Notably, the proportion of MSBm+ boutons in layer 4Cβ was twice that observed in the other two layers. This elevated presence of MSBm+ boutons in layer 4Cβ may reflect the higher density of geniculate afferents in this thalamorecipient layer (Hubel & Wiesel, 1972; Hendrickson et al. 1978). In our previous study using VGlut2 labeling, we found that layer 4Cβ had nearly double the density of TC afferents compared to 4Cα (Garcia-Marin et al. 2019).

These MSBm+ boutons are particularly relevant because they are the primary conduits of feedforward visual information. While earlier studies reported that thalamocortical (TC) input accounts for only 5–8% of total synapses in layer 4 (Peters et al. 1994; Latawiec et al. 2000), more recent analyses in primate V1 (Garcia-Marin et al., 2019) and rodent S1 (Sadaka et al. 2003; Bopp et al. 2017) suggest TC input levels are 2–4 times higher than those earlier estimates.

Motivated by the differences observed in the proportion of bouton categories across layers, we conducted a more detailed morphological analysis of the axonal profiles to determine whether structural differences exist between categories. This analysis is relevant to cortical circuitry, as MSBm+—and potentially some SSBm+—boutons in layer 4C are likely to correspond to thalamocortical terminals originating from the lateral geniculate nucleus (LGN), which are labeled with VGlut2 (Garcia-Marin et al. 2019). In contrast, the predominant SSBm− boutons are more likely to represent recurrent intracortical inputs across all layers and are typically associated with VGlut1 expression (Alonso-Nanclares et al. 2004). Notably, a substantial fraction of boutons in both the MSBm+ and SSBm+ categories contain mitochondria, which may reflect higher energetic or functional demands. Identifying structural differences across bouton categories and layers may provide insight into their distinct functional roles.

### Bouton Volume and Presynaptic Specialization

A major distinction between bouton categories was their volume. MSBm+ boutons - associated with thalamocortical (TC) inputs in layer 4C-were 2–4 times larger than SSBm+ boutons and 5–12 times larger than SSBm− boutons. These findings align with previous studies showing that the areas of the LGN-derived boutons are approximately three times greater than intracortical boutons in ferret V1 (Nahmani & Erisir, 2005; Erisir & Dreusicke, 2005). In our dataset, MSBm+ boutons in layer 4C were also about three times larger than those in layer 3B, consistent with values reported by Hsu et al. (2017), who found a mean volume of 0.45 μm³ for multisynaptic boutons in macaque layer 3. Such large bouton areas (Nahmani & Erisir, 2005) and volumes (Rodríguez-Moreno et al. 2018; Bopp et al. 2017) are a hallmark of TC terminals in primary sensory cortical areas. These TC boutons have been shown to express VGlut2 in ferret (Nahmani & Erisir, 2005) and the distributions of VGlut2 puncta in non-human primate V1 (Balaram et al. 2014; Garcia-Marin et al. 2013, 2019), and human V1 (Garcia-Marin et al. 2013) are consistent with the TC terminal organization. Notably, the volumes of MSBm+ boutons in macaque layer 4C (Fig. 7E; Table 7) were considerably larger than those reported for VGlut2+ boutons in mouse barrel and somatosensory cortex (0.306–0.45 μm³; Bopp et al. 2017; Rodriguez-Moreno et al. 2018), suggesting that the size of thalamocortical driver terminals increases across the phylogenetic scale—potentially reflecting greater synaptic efficacy or computational complexity in higher-order species.

In general, larger bouton size may be functionally linked to higher energetic requirements. The mitochondria occupied 20% of the total bouton volume, across the two categories with mitochondria (MSBm+ and SSBm+), independent of layer (Fig. 8). This result is consistent with previous findings in TC terminals (Bopp et al. 2017; Rodríguez-Moreno et al. 2018). The mitochondria are essential for critical processes involved in the neurotransmission by supplying ATP - required for neurotransmitter uptake and vesicle recycling, and by buffering calcium (David & Barrett, 2000; Kuromi & Kidokoro, 2005; Rizzoli & Betz, 2005; Vos et al. 2010; Denker & Rizzoli, 2010). TC terminals display a high release probability that ensures efficient, reliable signal transmission, particularly during high-frequency stimulation (Gil et al. 1999; Lee & Sherman, 2008; Saez & Friedlander, 2009). The co-occurrence of large bouton volume and high mitochondrial content in MSBm+ profiles thus likely reflects the functional demands placed on TC synapses in primate cortex.

Ultrastructural studies in the cerebral cortex, cerebellum, and hippocampus have shown that multiple presynaptic and postsynaptic features are positively correlated. Specifically, bouton volume, the number of synaptic vesicles, active zone size, PSD area, and spine head volume tend to scale together (Harris & Stevens, 1988; Schikorski & Stevens, 1997, 1999). In our dataset, the expanded volume of MSBm+ boutons may facilitate greater synaptic output by allowing for a larger pool of synaptic vesicles. This interpretation is consistent with prior findings that bouton volume correlates with vesicle number across species and cortical region (Pierce & Mendell, 1993; Germuska et al. 2006;

Zikopoulos & Barbas, 2007; Schmuhl-Giesen et al., 2022). For example, it has been reported that thalamocortical boutons in the prefrontal cortex show a positive relationship between bouton size and vesicle content, with large boutons containing over twice as many vesicles as small ones (Zikopoulos & Barbas, 2007). Overall many studies indicate that bouton size may correlate with synaptic strength because larger boutons have an increased size of their readily releasable vesicle pool (Tong & Jahr, 1994; Murthy et al. 1997, 2001; Schikorski & Stevens, 1999; Li et al. 2007. Although we did not quantify vesicle pools, the larger size of the MSBm+ boutons may support greater synaptic output and higher release rates, consistent with their morphology and presumed functional role because of a larger vesicle reservoir.

Consistent with this structural scaling, the largest MSBm+ terminals in our current study were those forming the most synapses—up to six in one single bouton. The presence of multisynaptic boutons in thalamocortical terminals has been reported across species and sensory systems, with individual boutons forming 3–8 active zones (Hamos et al. 1987; Bopp et al. 2017). In contrast, the smallest boutons (SSBm−) consistently formed only a single synapse, underscoring the structural and likely functional diversity among bouton subtypes. Together, these findings indicate that bouton volume serves as a reliable anatomical proxy for synaptic strength and output potential in the primate brain.

### Postsynaptic Features and Target Specificity

In a similar way that the presynaptic parameters scaled with bouton volume, the postsynaptic characteristics, such as PSD surface area, the presence of PSD perforations, and spine head volume, scale with the size of the AMPA receptor population (Desmond & Weinberg, 1998; Matsuzaki et al. 2001; Ganeshina et al., 2004; reviewed in Bourne & Harris, 2008). Larger or perforated PSDs and bigger spine heads mark synapses with more receptors, larger quantal amplitudes, and often higher release probability—core elements of greater synaptic strength.

In the current study the frequency distribution analysis showed that PSD’s area from MSBm+ and SSBm-were larger in the thalamo-recipient layers compared to layer 3. Previous data have reported the that PSD area varies in different areas and species. The PSD area of Vglut2+ terminals were larger than in Vglut2-terminals, in layer 4 of mouse M1 and S1 (Vglut2+: 0.064 and 0.042 µm² M1 and mouse S1; Vglut2-: 0.056 and 0.039 µm² M1 and S1, respectively) (Bopp et al. 2017). In primates, PSD area in layer 3 of macaque V1 and LPFC, were 0.082 and 0.116 µm², respectively) (Medalla & Luebke, 2015). These results suggest an increase in the PSD area on the phylogenetic scale and in relation to the bouton physiological properties. Our layer-3 PSD area values fall between those previously reported for macaque V1 and LPFC (Medalla & Luebke, 2015) and expand the results demonstrating that the PSD areas in the TC recipient layers are larger than in the non-TC recipient layers. The current data also confirmed Bopp’s results as our MSBm+, that probably represent their Vgtlut2+ terminals-area larger than the SSB -presumably Vglut2-).

Our current study also demonstrate that both bouton volume and PSD area vary by different categories and layers. If the synaptic strength is related to the ultrastructural morphology, we assume that there will be a positive correlation between bouton volume and PSD area. Previous studies has demonstrated that there is a positive linear relation between bouton volume and PSD area irrespective of the neuronal type or species examined (Pierce & Lewin, 1994; Bopp et al. 2017; Hsu et al. 2017). Our data demonstrated that for every single bouton category there is an positive correlation between bouton volume and the sum of the PSD area, reflecting a potential increase in the synaptic strength in the larger bouton.

In the current study we identified the postsynaptic target of each bouton. Each asymmetric synapse was classified as either on dendritic spines (axospinous) or on dendritic shafts (axodendritic). As expected, the majority of asymmetric synapses contacted dendritic spines—74% in layer 3B and 65% in layer 4C (see Table 4)—consistent with prior studies in cat, rat, and rhesus V1 (Beaulieu & Colonnier, 1985; Peters & Feldman, 1976; Medalla & Luebke, 2015; Santuy et al. 2018). These values further support the general principle that excitatory synapses preferentially target spines of pyramidal or spiny stellate neurons (Anderson et al. 1994; Arellano et al. 2007). To further explore this trend, we also analyzed postsynaptic target preference across bouton categories. MSBm+ boutons showed the strongest preference for spines, with 87% of their synapses being axospinous, across layers, compared to 66–67% for SSBm and SSBm+ boutons. This pattern supports the idea that MSBm+ boutons—likely of thalamic origin—preferentially target excitatory neurons reinforcing the functional distinction between TC and IC inputs.

Finally, across primate cortex the shape of the postsynaptic density (PSD) is also a structural marker of synaptic strength. Our 3-D reconstructions show that, independent of bouton category, macular junctions dominate in macaque V1 layers 3B, 4Cα and 4Cβ (92 %, Table 5). When bouton category is factored into the comparison (Table 6), botouns associated with complex PSDs (horseshoe and perforated) are more likely to bethe boutons with mitochondriathat also have the largest volumes and putatively larger vesicle pools. This result is consistent with those of Medalla & Luebke (2015) who showed that perforated PSDs are over-represented on the large, vesicle-rich boutons of macaque LPFC. In studies of human temporal-lobe Yakoubi et al. (2019a,b) and Schmuhl-Giesen et al. (2022) also reported that horseshoe and perforated PSDs, although a minority overall, were more frequently localized with larger boutons. Together with our V1 data, the enrichment of horseshoe/perforated PSDs associated with larger boutons suggests this is a common feature across many cortical regions in primates.

### Functional implications

Our study demonstrates layer-specific specialization in synaptic architecture, with clear differences between TC recipient layers (4Cα and 4Cβ) and non-TC recipient layers (3B). TC synapses are characterized by high efficacy, rapid transmission, and reliable activation, ensuring that the primary visual signal is faithfully relayed through the LGN to V1. In contrast, modulatory IC inputs support contextual modulation, gain control, and integration across receptive fields.

Electrophysiological studies have demonstrated that TC synapses are functionally stronger than their IC counterparts. In cat V1, TC synapses produce larger amplitude EPSPs, exhibit lower variability, and show strong short-term depression upon repeated stimulation (Stratford et al. 1996). Moreover, TC inputs have a higher innervation ratio and release probability, making them approximately 4.8 times more effective per connection than IC inputs, despite having similar quantal sizes (Gil et al. 1999).

Our anatomical findings support and extend these physiological results. In layer 4C, we found more MSBm+ boutons — which likely correspond to TC terminals —, which are up to 12 times larger than SSBm− boutons (presumably IC), form approximately three times more synapses, contain larger PSDs, and house a larger mitochondrial volume. These structural characteristics support a role for MSBm+ boutons in fast, high-fidelity transmission, and greater synaptic strength potential. In particular, mitochondrial enrichment may be critical for maintaining synaptic efficacy under sustained or high-frequency activation, due to its role in ATP production, calcium buffering, and vesicle mobilization (Jahn et al. 2003; Schneggenburger & Neher, 2005; Südhof & Rothman, 2009).

In conclusion, the neuropil architecture of layers 3B, 4Cα, and 4Cβ in macaque V1 reveals the structural basis of their functional specialization. The pronounced structural differences in synaptic boutons — particularly the distinctive features of MSBm+ terminals — are consistent with the known roles of TC and IC inputs and underscore the importance of synaptic morphology in shaping cortical information processing.

## Acknowledgements

Some of this work was performed at the Simons Electron Microscopy Center located at the New York Structural Biology Center, supported by the NIH Common Fund Transformative High Resolution Cryo-Electron Microscopy program (U24 GM129539), and by grants from the Simons Foundation (SF349247) and NY State Assembly. We thank Ed Eng, Ashleigh Raczkowski and William Rice for microscope assistance with data collection. This work was also supported by NIH grants: NIH Grant SC3NS127766 (VGM), PSC-CUNY rounds 52, 53, 55 (VGM), and P30EY013079 (MJH).

---

**Supplementary Figure 1.**
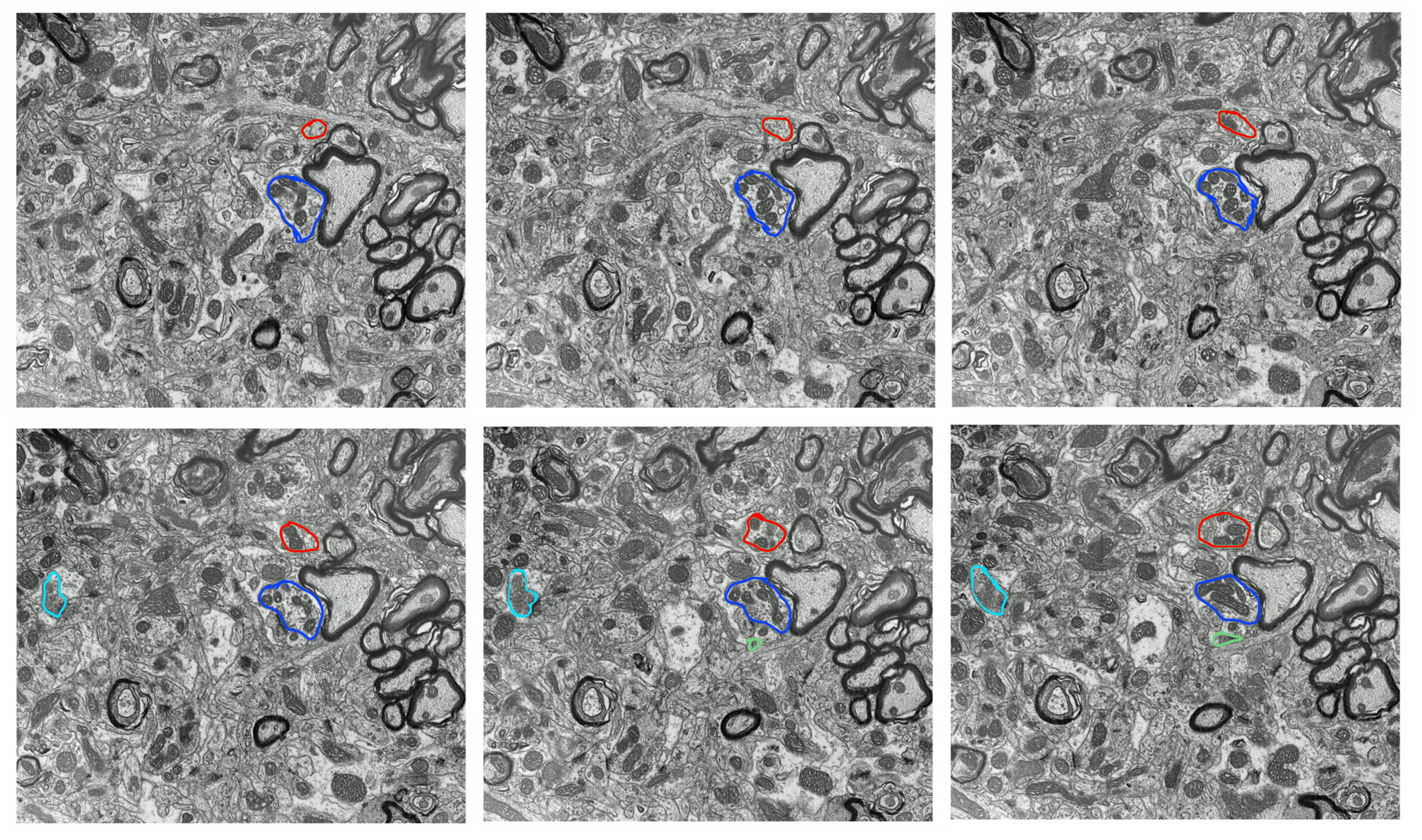
: Serial FIB-SEM images from layer 4Cβ of macaque V1 illustrating terminal boutons reconstructed in AMIRA. Six consecutive sections are shown (z-step: 60 nm), with each panel showing one example of each bouton category: MSBm+ (blue), SSBm+ (red), MSBm- (cyan), and SSBm- (green). Boutons are outlined in color to highlight their identity and continuity across sections.

**Supplementary Figure 2:**
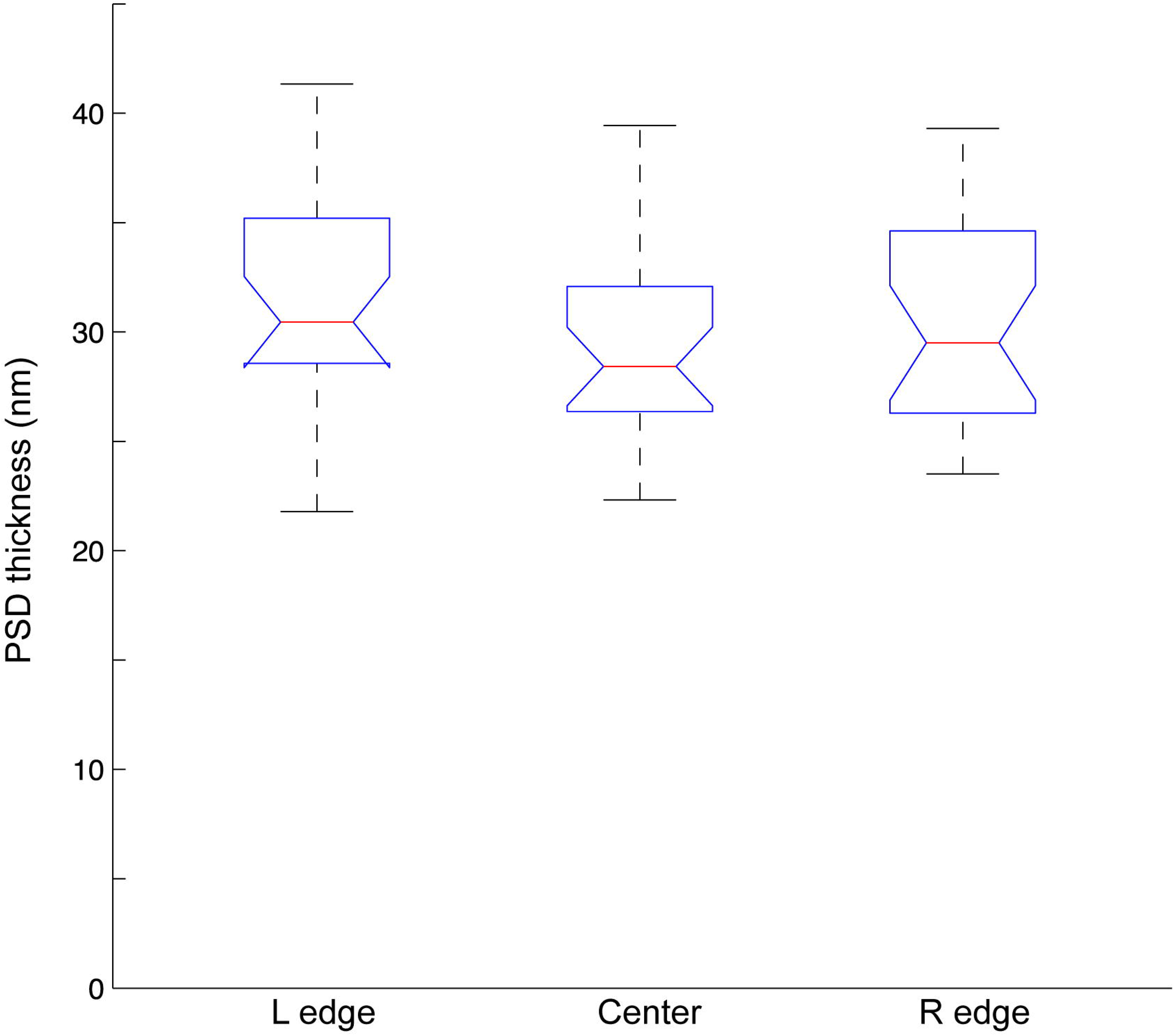
Thickness of the PSD depth measured at different locations of the PSD, on the left, the right, and in the center. No significant difference was found in the thickness of the PSD at the different locations (KW, p=0.256, ns). Each box spans the interquartile range (IQR), from the 25th percentile (bottom edge) to the 75th percentile (top edge). The red line inside the box indicates the median, and the notch represents a confidence interval around the median. Whiskers extend to the most extreme data points within 1.Sx IQR from the box.

**Supplementary Figure 3:**
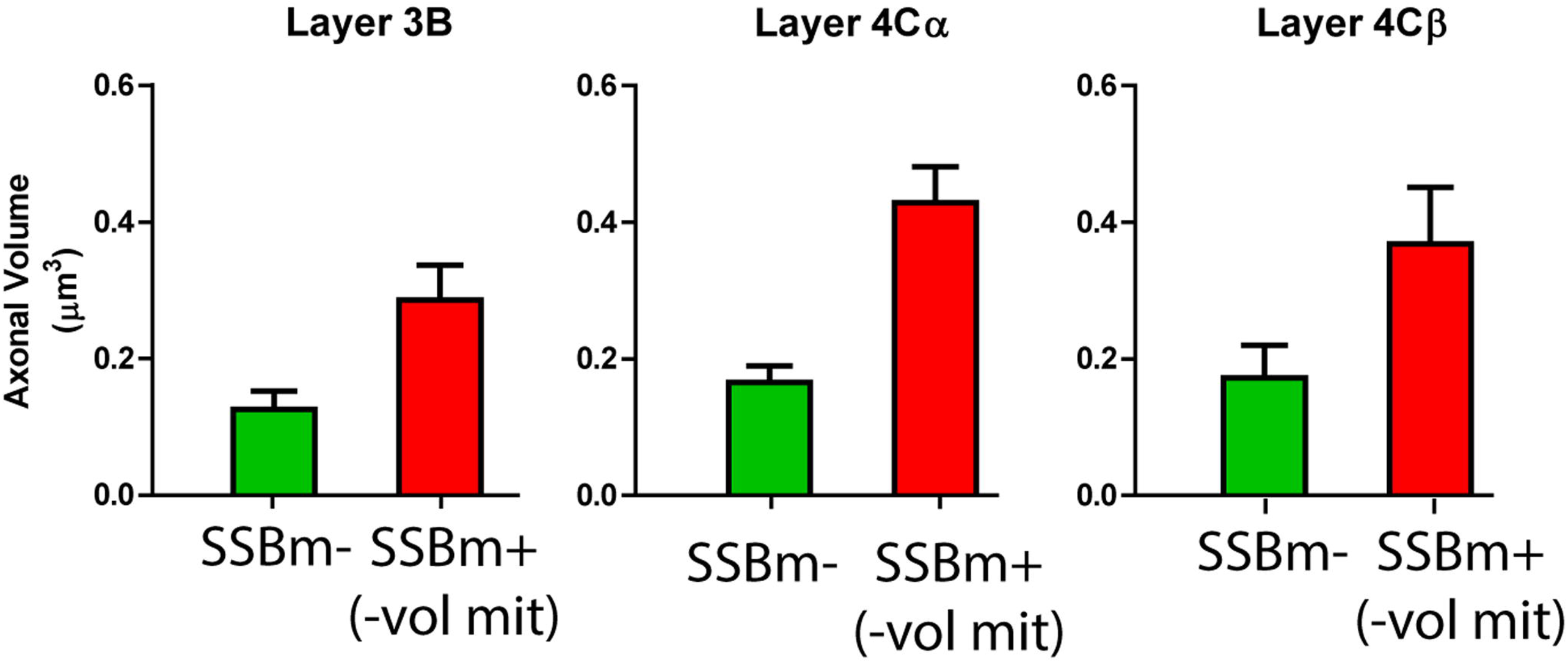
Comparison of the volume of SSBm-terminals with the volume of SSBm+ terminals, subtracting the mitochondrial volume. The results demonstrate that the bouton volume, excluding mitochondria, of the SSBm+ terminals was significantly larger than that of the SSBmterminals across all three layers suggesting that the increase in volume of the SSBm+ terminals cannot be solely attributed to the volume of the mitochondria (Layer 3B: Mann-Whitney test, ***; Layer 4Cα: Mann-Whitney test,***); Layer 4CP: Mann-Whitney test,**).**** p < 0.0001; *** p < 0.001; ** p < 0.01; * p < 0.05; ns p > 0.05

