## Supplementary Table 3 for "Three-dimensional ultrastructural differences between thalamic and non-thalamic recipient layers in macaque V1"

**Suppl. Table 3:** Density of synapses estimated in layer 4Cβ using 3D EM (synapses/μm^3^):

| **Animal** | **volume (μm^3^)** | **total synapses** | **Total synaptic**  **Density**  (synapses/μm^3^) | **Asymmetric synaptic density**  (synapses/μm^3^) | **Symmetric synaptic density**  (synapses/μm^3^) |
| --- | --- | --- | --- | --- | --- |
| M617 | 54.76 | 44 | 0.804 | 0.699 | 0.105 |
| M617 | 46.46 | 40 | 0.861 | 0.749 | 0.112 |
| M617 | 34.91 | 29 | 0.706 | 0.614 | 0.092 |
| M617 | 41.06 | 26 | 0.633 | 0.551 | 0.082 |
| M645 | 104.69 | 93 | 0.888 | 0.773 | 0.115 |
| M645 | 125.29 | 123 | 0.982 | 0.854 | 0.128 |
| M645 | 110.39 | 64 | 0.580 | 0.505 | 0.075 |
| M645 | 87.01 | 41 | 0.471 | 0.410 | 0.061 |
| M645 | 122.33 | 129 | 1.055 | 0.918 | 0.137 |
| M645 | 128.43 | 118 | 0.919 | 0.800 | 0.119 |
| M645 | 86.65 | 43 | 0.496 | 0.432 | 0.064 |
| M645 | 103.88 | 93 | 0.895 | 0.779 | 0.116 |
| WW | 81.17 | 52 | 0.641 | 0.558 | 0.083 |
| WW | 71.49 | 31 | 0.434 | 0.378 | 0.056 |
| WW | 43.76 | 39 | 0.891 | 0.775 | 0.116 |
| WW | 200.18 | 141 | 0.704 | 0.612 | 0.092 |
| WW | 197.69 | 144 | 0.728 | 0.633 | 0.095 |
| WW | 68.77 | 50 | 0.720 | 0.626 | 0.094 |
| **Mean± SEM** | **94.886±11.360** | **72.667±9.915** | **0.751±0.042** | **0.653±0.037** | **0.098±0.005** |

The samples came from blocks from 3 animals. Volumes (shown in columns 2), that contained only neuropil (excluding blood vessels, somas or myelinated axons), were analyzed from each block. The average synaptic density from the neuropil only samples was calculated (Mean**±SEM**)
